# Structural and Functional Diversity among Agonist-Bound States of the GLP-1 Receptor

**DOI:** 10.1101/2021.02.24.432589

**Authors:** Brian P. Cary, Peishen Zhao, Tin T. Truong, Sarah J. Piper, Matthew J. Belousoff, Radostin Danev, Patrick M. Sexton, Denise Wootten, Samuel H. Gellman

## Abstract

Recent advances in G protein-coupled receptor (GPCR) structural elucidation have strengthened previous hypotheses that multi-dimensional signal propagation mediated by these receptors is, in part, dependent on their conformational mobility. However, the relationship between receptor function and static structures determined via crystallography or cryo-electron microscopy is not always clear. This study examines the contribution of peptide agonist conformational plasticity to activation of the glucagon-like peptide-1 receptor (GLP-1R), an important clinical target. We employ variants of the peptides GLP-1 and exendin-4 to explore the interplay between helical propensity near the agonist N-terminus and the ability to bind to and activate the receptor. Cryo-EM analysis of a complex involving an exendin-4 analogue, the GLP-1R and G_s_ protein revealed two receptor conformers with distinct modes of peptide-receptor engagement. Our functional and structural data suggest that receptor conformational dynamics associated with flexibility of the peptide N-terminal activation domain may be a key determinant of agonist efficacy.

---

G protein-coupled receptors (GPCRs) are critical conduits for intercellular communication. These membrane-embedded proteins transmit information borne by extracellular molecules to the cell interior. Signal transduction is mediated by agonist-facilitated conformational changes in the receptor that are sensed by intracellular transducers, such as G proteins and arrestins.^1^ Understanding mechanisms governing agonist activation of GPCRs is integral to interrogation of physiological processes controlled by these receptors and offers a basis for developing therapeutic agents. Recent methodological advances have provided molecular-level snapshots of GPCR structure, including ligand-induced changes in GPCR structure, and of interactions between GPCRs and intracellular partner proteins.^2^ Nevertheless, it is emerging that static structures determined via x-ray crystallography or cryo-electron microscopy (cryo-EM) cannot always be extrapolated to understand receptor and transducer activation, which are inherently dynamic processes.^3^ Here we describe an integrated chemical, pharmacological and structural approach to elucidate mechanisms of signal transduction by the glucagon-like peptide-1 receptor (GLP-1R), based on comparisons involving two natural agonists, GLP-1 and exendin-4, and rationally designed analogues of these peptides.

The GLP-1R is a class B1 peptide hormone GPCR that plays a critical role in glucose metabolism, and synthetic agonists of this receptor are used to treat type 2 diabetes and associated comorbidities.^4,5^ The primary endogenous GLP-1R agonist is the fully processed peptide, GLP-1(7-36)-NH_2_.^6^ Class B1 receptors feature a large extracellular domain (ECD) in addition to the ubiquitous heptahelical transmembrane domain (TMD). Initial agonist-receptor contact occurs between the C-terminal portion of the peptide and the ECD; the N-terminal portion of the agonist subsequently engages the TMD core, facilitating conformational changes that are registered by the G protein and other intracellular partners (Fig. 1a).^7–9^ Most agonist C-terminal regions are α-helical when bound to class B1 receptor ECDs,^10,11^ but insights into the structure of the agonist N-termini embedded in receptor TMDs have emerged only recently. For example, a co-crystal structure of the GLP-1R bound to a short GLP-1-derived agonist peptide and cryo-EM structures of this receptor complexed to a heterotrimeric G protein and bound to either the endogenous agonist, GLP-1, or a synthetic peptide, ExP5, have been reported.^12–15^ In each case, α-helical secondary structure extends to the TMD-engaged N-terminus of the bound peptide. Comparable observations were reported for agonist peptides bound to several other class B1 GPCRs.^16–25^ In contrast, structures of other class B1 GPCRs bound to calcitonin (CT),^26^ calcitonin gene related peptide (CGRP),^27^ maxadilan,^20^ corticotropin-releasing factor (CRF)^22^ or urocortin-1 (UCN1)^28^ reveal a loop secondary structure near the agonist N-terminus, followed by either a short (CT-family peptides) or extended α-helix. In each of these cases, helicity near the N-terminus is disfavored by sequence. For CT, maxadilan, and CGRP, a disulfide linkage precludes helix propagation to the N-terminus, while for UCN1 and CRF the presence of proline residues near the N-terminus discourages local α-helicity.

**Fig. 1.**
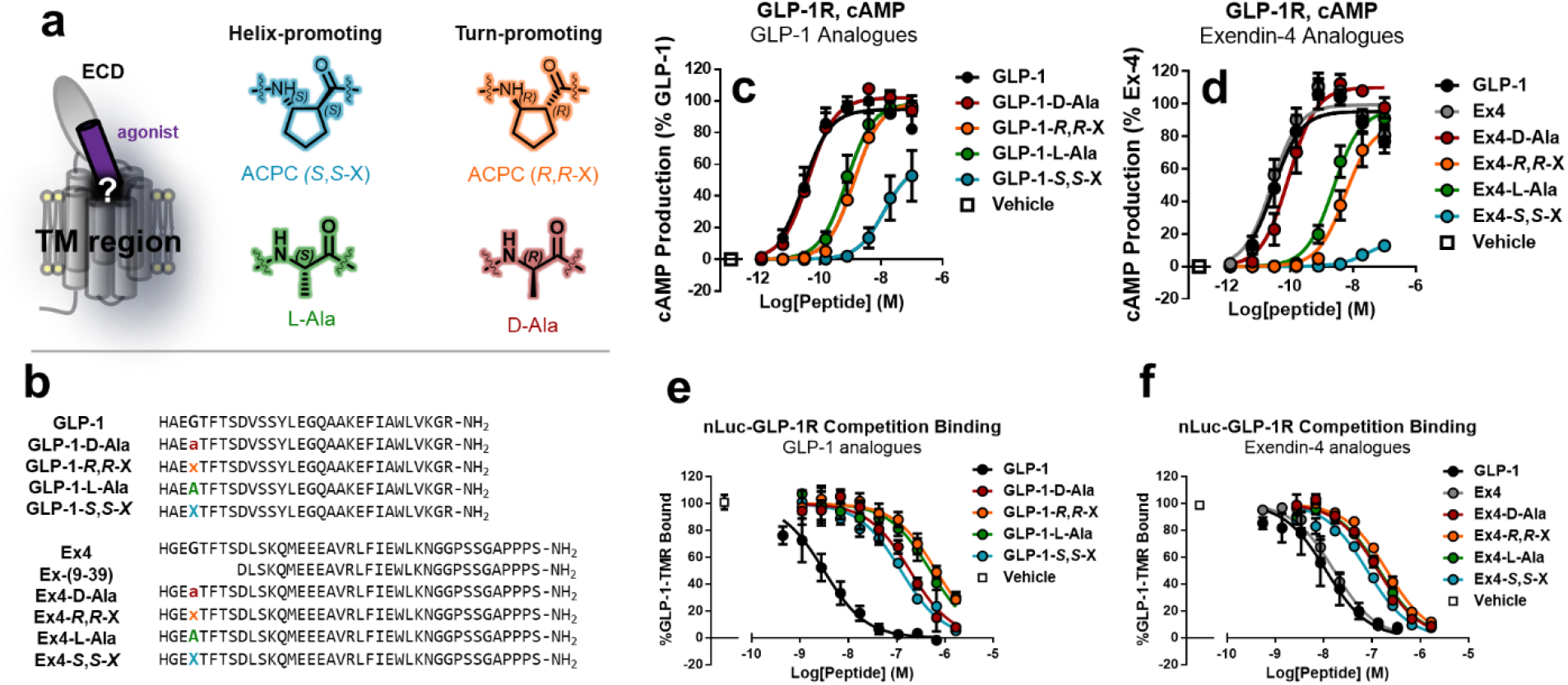
Probing the N-terminal structure of GLP-1 and Exendin-4 with single substitutions. **a,** Left: Cartoon depiction of an agonist peptide (purple) bound to a class-B GPCR. The extracellular domain (ECD) and transmembrane (TM) are labeled. Right: Amino acid residues used to probe the active state of GLP-1. **b**, Sequences of Exendin-4, GLP-1, and analogues. Lowercase ‘a’ represents D-Ala, uppercase ‘X’ represents (*S*,*S*)-X, and lowercase ‘x’ represents (*R*,*R*)-ACPC. **c**-**d**, Activation of GLP-1R-FLAG by GLP-1, Exendin-4, and analogues as measured by cAMP. Data points represent the mean of three independent experiments. **e-f**, Equilibrium Nluc-GLP-1R competition binding BRET assay performed with intact, NaN3-treated HEK293GS22 cells. Data points represent the mean of either three or four independent experiments, for **e** and **f** respectively. Error bars represent standard error.

The current work is predicated on previous suggestions that GLP-1 activity depends on adoption of a reverse turn near the peptide N-terminus, which raises the possibility that the N-terminal helical conformation in the cryo-EM structure of receptor-bound GLP-1^12^ may not fully capture structural requirements for signaling. This consideration is important because efforts to engineer therapeutic GLP-1R agonists might target a conformation that binds tightly to the signal-propagating form of the receptor. Evidence for an N-terminal reverse turn in GLP-1R agonists has emerged from NMR characterization of isolated peptides^29,30^ and modeling of peptide-receptor complexes.^31,32^ Bioinformatics analysis of multiple hormones, including GLP-1, predict a conserved helix-capping motif near the agonist N-terminus^33^ that would favor non-helical conformations in the segment preceding the cap. A survey of peptide-activated GPCRs led to speculation that turn-like structures might be a common motif among agonists^34^ and that N-terminal region flexibility may be important for class B1 peptide agonists.

Previous efforts to stabilize the proposed turn conformation near the GLP-1 N-terminus via side chain cross-linking produced mixed results. In some contexts, such cross-linking provided potent agonists, but other cross-linking efforts caused sharp declines in potency.^32,35^ Molecular modeling suggested that α-helical and β-turn conformations near the N-terminus should be energetically comparable for these cross-linked peptides, and that agonist potency was better correlated with computationally predicted α-helix propensity than with β-turn propensity.^35^ The β-turn hypothesis remains plausible, however, because of early studies with two diastereomeric analogues of GLP-1(7-36)-NH_2_, containing either Gly10→L-Ala or Gly10→D-Ala substitution. Watanabe et al. showed that the L-Ala diastereomer was nearly 100-fold less potent than GLP-1 itself in terms of stimulating insulin release from isolated rat pancreases, while the D-Ala diastereomer matched GLP-1 in potency.^36^ Gly is frequently found at the center of β-turn-forming segments, and Gly residues in such turns often display backbone torsion angles that are unfavorable for L-amino acid residues.^37,38^ Gly10 of GLP-1 is highly conserved across species.^39^ The observations of Watanabe et al. raise the possibility that signal transduction mediated by GLP-1 is promoted by the accessibility of a reverse turn centered on Gly10, rather than restriction to the α-helical secondary structure observed via cryo-EM for this region of receptor-bound GLP-1.^40^ Studies with peptide and protein model systems indicate that replacing Gly with L-Ala stabilizes a right-handed α-helical conformation by up to 1 kcal/mol,^41–43^ while replacing Gly with D-Ala *destabilizes* the α-helical conformation by up to 0.5 kcal/mol.^41^

## Results

### GLP-1R peptide agonists with D-Ala are more potent than those with L-Ala at the fourth position

We re-examined^36^ the agonist activity of Gly10→L-Ala and Gly10→D-Ala variants of GLP-1(7-36)-NH_2_ (Fig. 1b, 1c) with HEK293 cells transiently expressing the GLP-1R and stably expressing the GloSensor™ protein for detection of cAMP.^44^ Stimulation of intracellular cAMP production is typically used to monitor GPCR-modulated activation of the stimulatory G protein, G_αS_. Both Ala-containing diastereomers matched GLP-1 in terms of the maximum level of cAMP production. However, while the Gly10→D-Ala analogue was indistinguishable from the native hormone in terms of potency (EC_50_), the Gly10→L-Ala was ~24-fold less potent (Fig. 1c; Table 1). This behavior is qualitatively consistent with earlier observations.^36,45^ We explored the generality of these observations by evaluating Ala-containing derivatives of exendin-4 (exenatide) (Fig. 1b), a potent GLP-1R agonist isolated from a lizard venom that is used to treat type 2 diabetes.^46^ Exendin-4 and GLP-1 are very similar over the first 11 residues, and Gly10 of GLP-1 corresponds to Gly4 of exendin-4 (Fig. 1b). The Gly4→D-Ala variant of exendin-4 was only slightly less potent than exendin-4 itself in terms of cAMP production, but the Gly4→L-Ala variant was ~30-fold less potent, which parallels the trend among GLP-1 analogues (Fig. 1d; Table 1). These data support the conclusion that an ability to access non-helical conformations near the N-terminus correlates with higher GLP-1R agonist potency.

**Table 1.** EC_50_ values, maximal responses, and IC_50_ values from 3-parameter sigmoidal fits for concentration-response data in Fig 1. EC_50_ rel. indicates cAMP production potency relative to GLP-1 by the quotient (Peptide EC_50_) / (GLP-1 EC_50_). IC_50_ rel. indicates the affinity relative to GLP-1 by the quotient (Peptide IC_50_) / (GLP-1 IC_50_) [a] GLP-1 was averaged over 6 sets of independent experiments. Uncertainties are expressed as standard error of the mean.

|  | cAMP Production |  |  | EC<sub>50</sub> rel. | Whole-Cell Affinity |  |  |
| --- | --- | --- | --- | --- | --- | --- | --- |
|  | pEC<sub>50</sub> | EC<sub>50</sub> (nM) | % Max |  | pIC<sub>50</sub> | IC<sub>50</sub> (nM) | IC<sub>50</sub> rel. |
| **GLP-1<sup>a</sup>** | 10.5 ± 0.1 | 0.031 | 95 ± 4 | 1 | 8.27 ± 0.07 | 5.4 | 1 |
| **GLP-1-D-Ala** | 10.4 ± 0.09 | 0.044 | 102 ± 4 | 1.4 | 6.73 ± 0.05 | 187 | 35 |
| **GLP-1-R,R-X** | 8.87 ± 0.1 | 1.4 | 99 ± 5 | 45 | 6.19 ± 0.06 | 650 | 120 |
| **GLP-1-L-Ala** | 9.12 ± 0.1 | 0.76 | 99 ± 5 | 24 | 6.31 ± 0.06 | 500 | 93 |
| **GLP-1-S,S-X** | 7.91 ± 0.2 | 12 | 60 ± 10 | 400 | 6.91 ± 0.08 | 120 | 22 |
| **Ex-4** | 10.6 ± 0.1 | 0.026 | 99 ± 4 | 0.8 | 7.79 ± 0.04 | 16 | 3 |
| **Ex-4-D-Ala** | 10.0 ± 0.1 | 0.093 | 110 ± 4 | 3 | 6.88 ± 0.03 | 134 | 25 |
| **Ex-4-R,R-X** | 8.25 ± 0.1 | 5.6 | 87 ± 5 | 180 | 6.69 ± 0.03 | 200 | 37 |
| **Ex-4-L-Ala** | 8.62 ± 0.08 | 2.4 | 96 ± 4 | 77 | 6.84 ± 0.04 | 143 | 26 |
| **Ex-4-S,S-X** | 7.54 ± 0.1 | 29 | 16 ± 2 | 940 | 7.10 ± 0.04 | 80 | 15 |

### GLP-1R agonist analogues with turn-promoting β-amino acids are more active than those with helix-promoting β-amino acids at the fourth position

Non-traditional substitutions can yield GLP-1 analogues that show distinctive behavior and these might also provide insight into agonist conformation.^47–50^ To this end, we explored a second set of substitutions at the key Gly residue in GLP-1 and exendin-4. This experimental design was based on previous comparisons of the conformations and biological activities of conventional peptides (comprised entirely of α-amino acid residues) with the properties of analogues in which at least one α residue was replaced with a β-amino acid residue. Mixed-backbone peptides containing up to 25-33% β residues can adopt an α-helix-like secondary structure.^51^ The constrained β residue derived from *trans*-(*S,S*)-2-aminocyclopentanecarboxylic acid ((*S,S*)-ACPC) is comparable to L-Ala in stabilizing a right-handed α-helix-like conformation (Fig. 1a).^52^ We previously showed that GLP-1 analogues with multiple (*S*,*S*)-ACPC substitutions in the C-terminal region, which is α-helical when bound to the ECD, display substantial agonist activity.^53,54^ In contrast, (*R,R*)-ACPC (Fig. 1a) destabilizes a right-handed α-helix-like conformation by >1 kcal/mol relative to (*S,S*)-ACPC or L-Ala.^52^ As observed for D-Ala,^55^ (*R*,*R*)-ACPC can replace Gly to stabilize turn segments.^56^ These precedents led us to compare diastereomeric derivatives of GLP-1 and exendin-4 in which Gly10 or Gly4, respectively, was replaced by either (*S,S*)-ACPC or (*R,R*)-ACPC. Although the steric bulk of the (CH_2_)3 side chain might diminish activity relative to the natural GLP-1R agonists, these replacements should test the hypothesis that GLP-1R agonist activity is higher for ligands that can access non-helical conformations near the N-terminus, compared to those that cannot. This hypothesis predicts that the Gly→(*R,R*)-ACPC analogue should be more active than the diastereomer containing (*S,S*)-ACPC.

The relative activities among ACPC-containing analogues of the two natural GLP-1R agonists were consistent with predictions of our hypothesis: the Gly10→(*R,R*)-ACPC analogue of GLP-1 was ~9-fold more potent than the (*S,S*)-ACPC diastereomer, and the Gly4→(*R,R*)-ACPC analogue of exendin-4 was ~5-fold more potent than the (*S,S*)-ACPC diastereomer in eliciting cAMP production (Fig. 1c, 1d). Moreover, both analogues containing (*S*,*S*)-ACPC had reduced maximum cAMP production compared to their diastereomers. Nonetheless, in each case, even the more potent diastereomer was an inferior agonist relative to the all-α prototype, by ~45-fold in the GLP-1 series and ~220-fold in the exendin-4 series (Fig. 1c, 1d, Table 1). The patterns of relative activity among ACPC-containing analogues of GLP-1 and exendin-4 support the hypothesis that access to non-helical conformations near the agonist N-terminus is important for GLP-1R activation.

### Peptides with helix-promoting β residues at the fourth position show relatively high affinity among the modified analogues

An agonist’s potency is influenced by both affinity for the receptor and the ability to shift the receptor into active conformations that transduce the signal via interaction with intracellular proteins.^57^ Agonist affinity for GPCRs has typically been measured via competition with a labelled probe ligand.^58^ We developed a competition assay based on detection of probe binding via bioluminescence resonance energy transfer (BRET). Key components for this assay were a version of human GLP-1R with the bright, bioluminescent protein NanoLuc (NLuc)^59^ fused to the N-terminus, and a GLP-1(7-36) derivative bearing a tetramethylrhodamine moiety linked to a lysine side chain at position 36. This assay can be performed without washing, providing advantages over conventional binding assays.^60^ Normalized IC_50_ values (relative to GLP-1) derived from this competition BRET assay, in intact cells at equilibrium, show that all four modifications at Gly10 of GLP-1 and all four modifications at Gly4 of exendin-4 cause substantial declines in affinity for the GLP-1R relative to the natural agonist. Effects of the substitutions on affinity were distinct from the effects of the substitutions on peptide potency, with the modified analogues for each peptide displaying relatively similar affinities to each other, but vastly different potencies for cAMP production (Fig. 1e, 1f; Table 1). For example, the Gly10→D-Ala analogue of GLP-1 is indistinguishable from GLP-1 itself in terms of EC_50_, but the analogue shows a ~35-fold diminution in affinity. This D-Ala analogue is ~17-fold more potent than the L-Ala diastereomer but binds only ~3-fold more tightly to the GLP-1R (Fig. 2a, 2b).

**Fig. 2.**
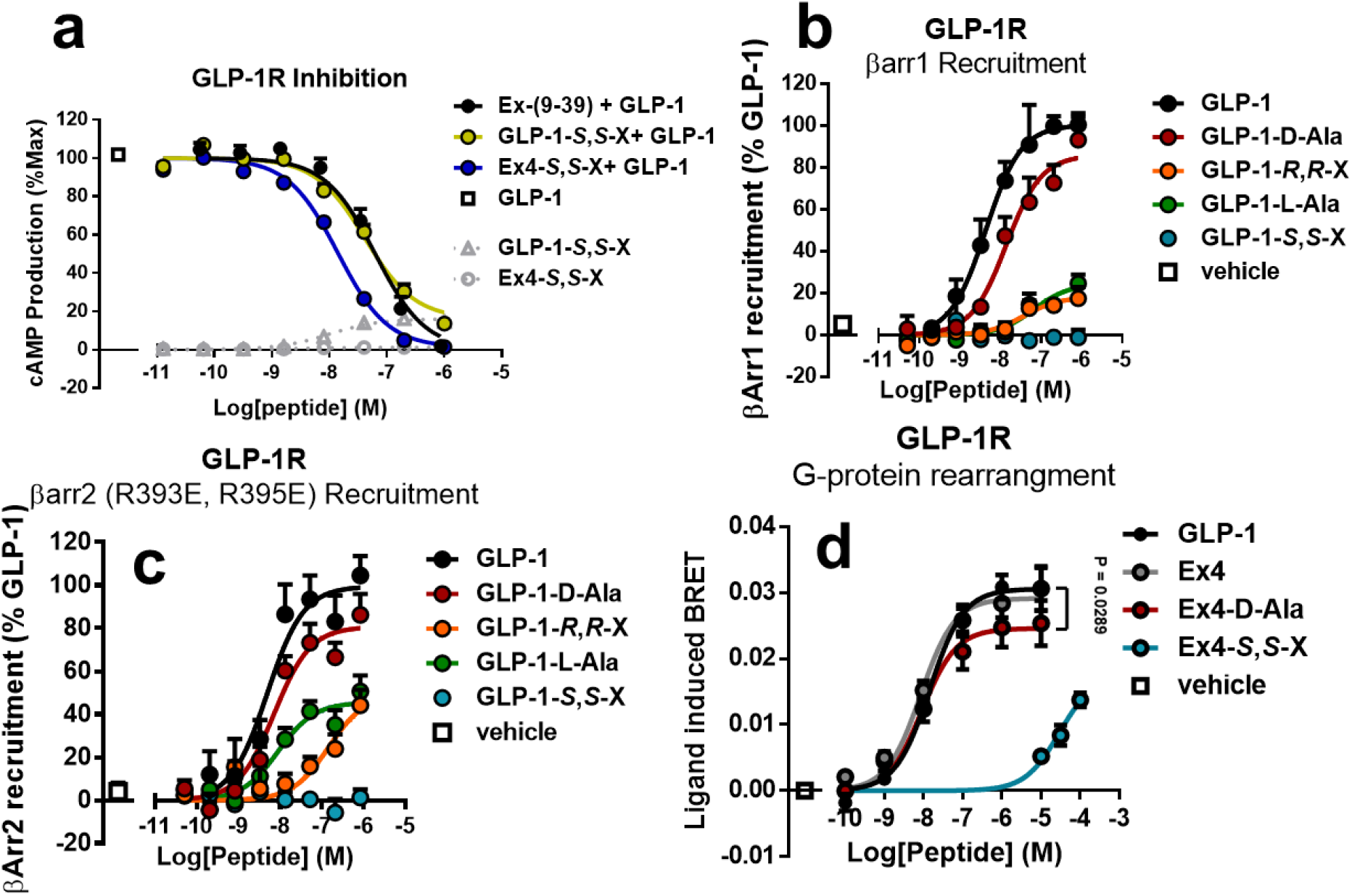
Further characterization of N-terminally substituted analogues. **a**, Inhibition of GLP-1 stimulated cAMP production in HEK293GS22 cells expressing hGLP-1R. Cells were preincubated for 15 min with increasing concentrations of ACPC substituted peptides or Ex (9-39) followed by stimulation with 0.25 nM GLP-1. Grey symbols with dotted connecting lines represent the cAMP accumulation in response to GLP-1 (*S*,*S*-X) and Ex4 (*S*,*S*-X) before addition of GLP-1. **b**, β-arrestin-1 recruitment to GLP-1R-Rluc8. **c**, β-arrestin-2 (R939E, R395E) recruitment to GLP-1R-Rluc8. **d**, Dose-response G-protein conformational rearrangement as measured by BRET between Gαs–nanoluc, Gβ_1_γ_2_–venus at a terminal timepoint (12 min). The P-value compares the fitted maximal responses of GLP-1 and Ex4-D-Ala. The P value was determined by one-way ANOVA with Bonferroni’s post-test. Data points represent the mean of three independent experiments. Error bars represent standard error.

### Ex-4-*S*,*S*-X potently inhibits GLP-1 mediated cAMP formation at GLP-1R

Among the four GLP-1 analogues, Gly10→(*S,S*)-ACPC is the least efficacious agonist but has highest affinity for the receptor. This affinity pattern is qualitatively paralleled among the four exendin-4 variants. We found that both (*S,S*)-ACPC-containing analogues could function as antagonists of GLP-1-induced cAMP production in HEK293GS22 cells. Indeed, the Gly4→(*S,S*)-ACPC derivative of exendin-4 proved to be an even more potent antagonist than exendin-(9-39), which is currently in clinical trials for the treatment of post-bariatric hypoglycemia (Fig. 2a, Table S1).^61^ Exendin-(9-39) cannot activate the GLP-1R because this peptide lacks N-terminal residues that engage the TMD core.^36^ The superior antagonist activity of the Gly4→(*S,S*)-ACPC derivative relative to exendin-(9-39) suggests that the N-terminus of the (*S,S*)-ACPC-containing peptide engages the TMD in a manner that is energetically favorable but ineffective for GLP-1R activation.

The relatively high affinity displayed by the Gly4→(*S,S*)-ACPC derivative of exendin-4 for the GLP-1R suggests that stable receptor-ligand complexes occur when helical secondary structure extends to the ligand N-terminus, consistent with the consensus conformation of GLP-1R peptide agonists in recent crystal and cryo-EM structures.^14,40^ However, the potency data for stimulation of cAMP production collectively suggest that signal transduction via G_αs_ is facilitated if the receptor-bound ligand retains flexibility and can access a non-helical conformation near the N-terminus. Assays of recruitment of β-arrestin-1 or −2 to the GLP-1R indicated a similar requirement for agonist N-terminal flexibility (Fig. 2b, 2c; Table S2). Further support for the functional importance of non-helical conformations near the agonist N-terminus was obtained from a BRET-based assay that monitors receptor-mediated changes in G protein conformation.^15,62^ The Gly4→D-Ala derivative of exendin-4 had similar potency to exendin-4 itself in inducing conformational changes in the heterotrimeric G protein, albeit with modestly lower maximal response, while the Gly4→(*S,S*)-ACPC derivative of exendin-4 was markedly less potent (Fig. 2d, Table S3).

### A cryo-EM structure of Ex4-D-Ala bound to GLP-1R/Gαs

To investigate the receptor-bound conformation of the potent exendin-4 analogue containing D-Ala in place of Gly4 (referred to below as Ex4-D-Ala), we undertook cryo-EM studies of the complex formed by this agonist with the GLP-1R.^63^ We co-expressed the human GLP-1R, dominant negative G_α s_,^64^ G_β1_, and G_γ2_ in *Trichoplusia ni* cells. Nanobody 35, excess peptide ligand (10 μM), and apyrase were added to form a complex, which was solubilized in lauryl maltose neopentyl glycol (LMNG)/cholesterol hemisuccinate (CHS) mixed-micelles as previously reported.^15,65,66^ This complex was purified by sequential anti-FLAG affinity and size exclusion chromatography in the presence of saturating ligand (2.5 μM) to yield a monophasic peak on SEC containing each of the components of the complex, which was confirmed in negative stain TEM (Fig. S3A-S3E). Although we were also able to form a GLP-1R/G-protein complex with the Gly4→(*S,S*)-ACPC analogue of exendin-4 (Fig. S3F-S3J), yields were poor, and the sample was too heterogenous by size-exclusion chromatography (Fig. S3G) and negative stain TEM (Fig. S3J) to warrant imaging by cryo-EM.

The purified Ex4-D-Ala complex was vitrified, and single particles were imaged on a Titan Krios TEM.^67^ After 2D and 3D classification of particle images, a consensus map with a nominal global resolution of 2.3 Å was resolved (Fig. S4, S5A). Despite some orientation bias (Fig. S5C), high local resolution in the receptor core and G protein enabled modeling of most of the complex including the N-terminus of the peptide within the receptor core (Fig. 3a, 4b, 4c, Fig. S5E). The local resolution in the ECD was lower but allowed for fitting of the ECD backbone and modelling of side chains in the peptide vicinity. Poor resolution was observed for the G_αs_ alpha-helical domain and for ICL3 (residues 59-204 and 338-340, respectively), so these segments were omitted from the atomic model.

**Fig. 3.**
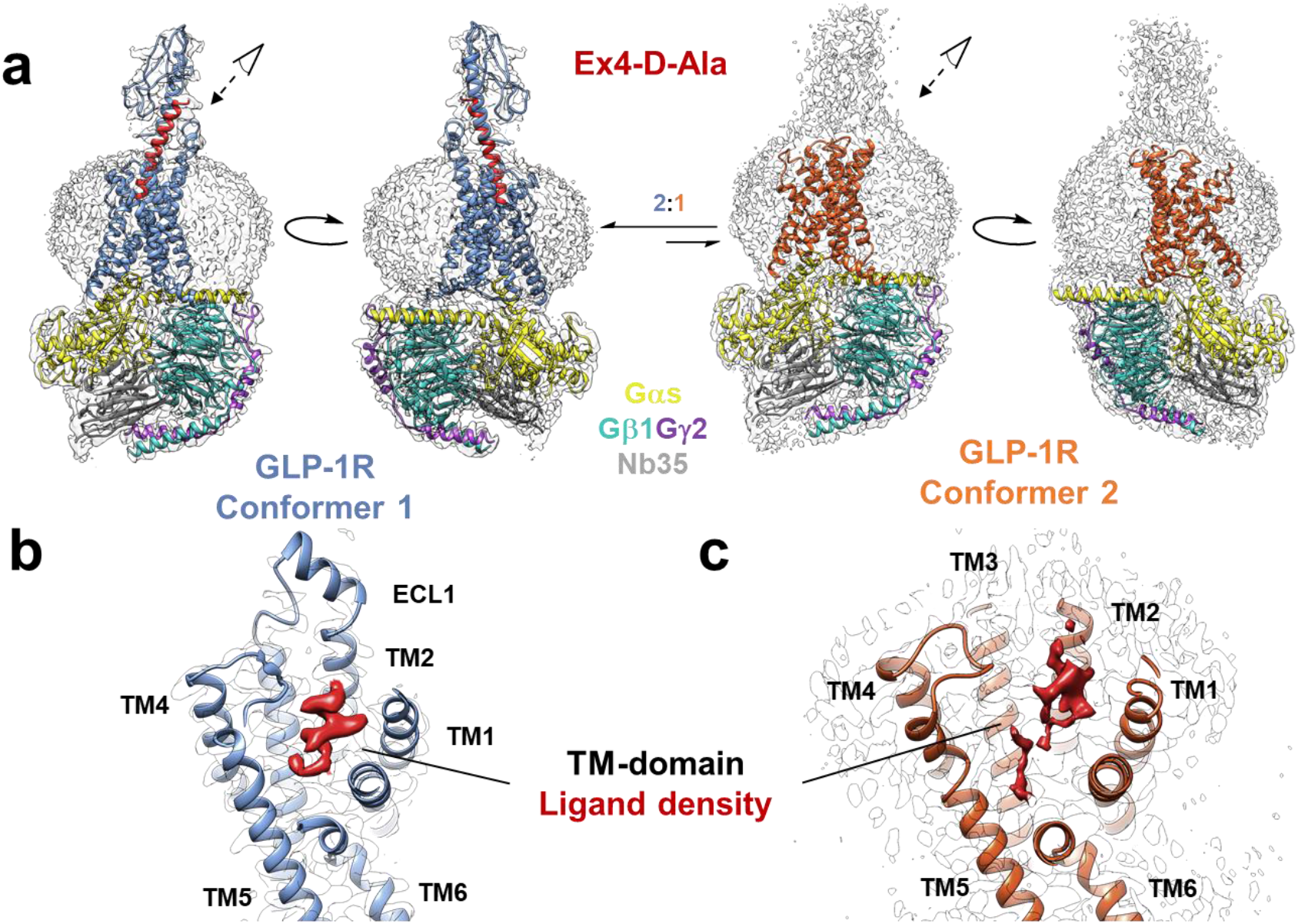
Cryo-EM structure of Ex4-D-Ala bound GLP-1R in complex with the heterotrimeric G-protein and nanobody 35. **a,** The models of the two conformers are shown within the cryo-EM derived density maps which are depicted as a transparent surface. GLP-1R in conformer 1 is colored blue, while GLP-1R in conformer 2 is colored orange. The number of particles used in the reconstruction indicated an approximately 2:1 ratio of Conformer 1 to Conformer 2. Dominant negative Gαs, Gβ1, Gγ, and nanobody 35 are colored yellow, aqua, purple, and gray, respectively. **b**, The orthosteric binding pocket of GLP-1R in conformer 1 is shown with the ECD and ECL3 removed for clarity. Ligand density is shown in red. **c**, The orthosteric binding pocket of GLP-1R in conformer 2 with ECL3 removed for clarity. The ligand density is shown in red.

**Fig. 4.**
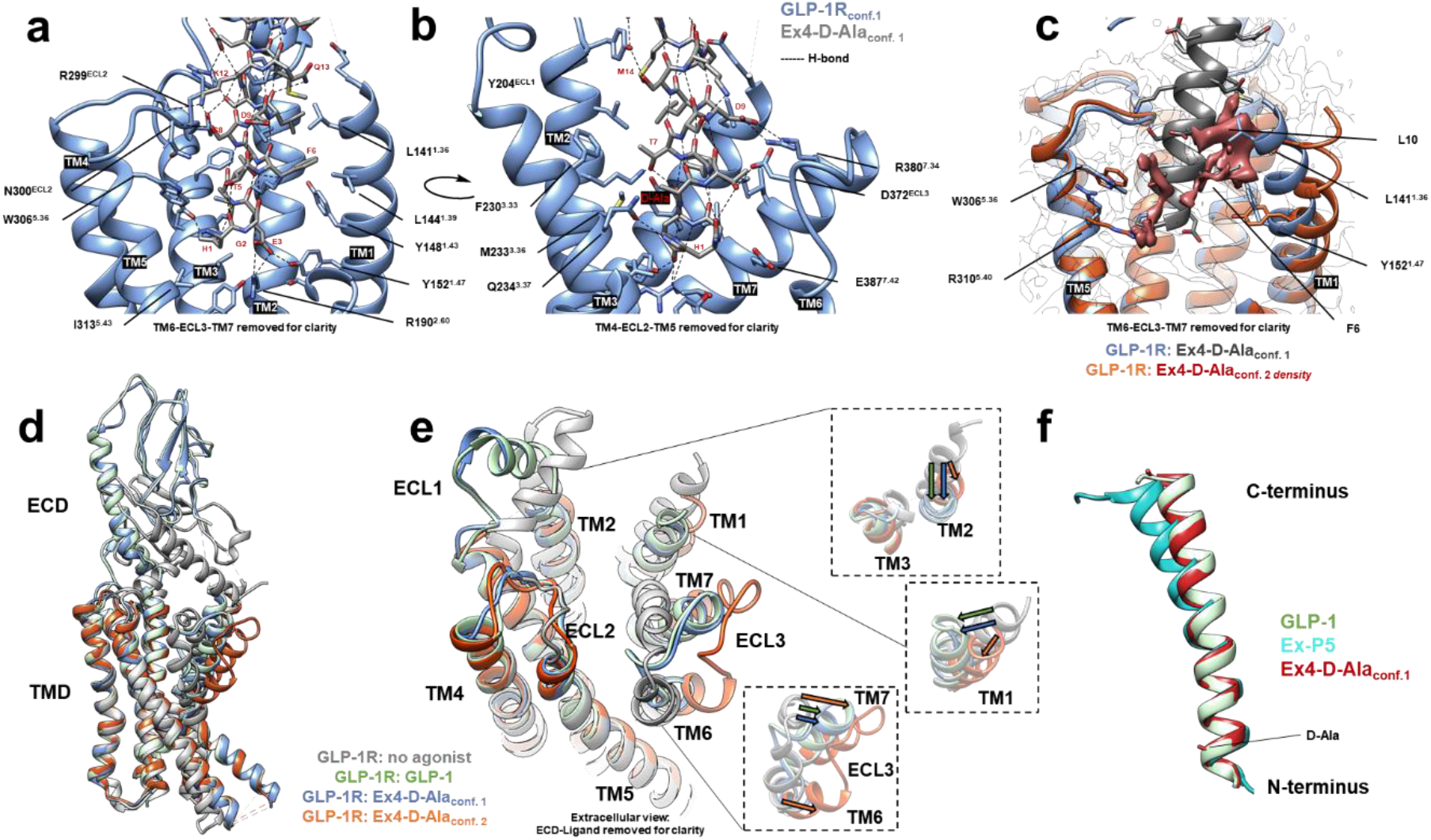
The structure of Ex4-D-Ala bound to GLP-1R. **a**, A close-up, side-view of the orthosteric binding pocket as determined in conformer 1. TM6, ECL3, and TM7 were removed for clarity. **b**, A close-up, side-view of the orthosteric binding pocket as determined in conformer 1 rotated relative to the view in Figure 5A. TM4, ECL3, and TM5 were removed for clarity. **c**, An overlay of conformer 1 and conformer 2 states of GLP-1R shown in blue and orange, respectively. Conformer 1 ligand is shown in grey and conformer 2 orthosteric density is colored red. **d**, A comparison of GLP-1 bound GLP-1R, GLP-1R without agonist or G-protein bound, and Ex4-(D-Ala) bound GLP-1R as observed in conformer 1 and 2 (colored blue and orange, respectively). **e**, An extracellular view of models compared in Fig. 4d, but with the extracellular domain removed for clarity. The boxes show movements of the structures relative to the no-agonist GLP-1R crystal structure. **f,** A comparison of positioning and conformation of three peptide agonists (GLP-1 in green, ExP5 in teal, and Ex4-D-Ala as observed in conformer 1 in red) when the receptor-G-protein complexes are aligned.

The Ex4-D-Ala complex is very similar to our recently published high-resolution structure of GLP-1 bound to the human GLP-1R (Fig. 4d).^12^ Even though Ex4-D-Ala shares the same C-terminal residues as the G protein-biased agonist exendin P5 (ExP5),^15,68^ Ex4-D-Ala adopted a distinct position relative to the receptor from that displayed by ExP5 (Fig. 4f, S9C), and Ex4-D-Ala induced an ECL1 conformation closer to that in the GLP-1-bound structure (Fig. 4e) than that in the ExP5-bound structure (Fig. S9D). In the consensus map, the agonist adopted an α-helical conformation along its entire length, despite the presence of D-Ala near the N-terminus (Fig. 4f). The D-Ala residue displayed right-handed helical Φ and Ψ torsion angles of −61° and −50°, respectively. The methyl side chain of D-Ala is close to the side chains of two receptor residues, M233^3.36^ and Q234^3.37^ (Fig. 4a, 4b, S8A), that influence the affinity and potency of the natural agonist GLP-1.^69^ Interactions of these receptor side chains with the agonist D-Ala side chain might compensate for helix-destabilizing effects of the D-Ala residue.

### Two distinct conformers were apparent within the cryo-EM dataset

The low resolution of the ECD led us to perform additional 3D classification of the particles from the consensus map (Fig. S4), which revealed a second conformation of the peptide-occupied receptor (conformer 2). Conformer 2 represented approximately one-third of the particles, with the remainder corresponding to the consensus conformation described above (conformer 1). Conformer 2 was refined to a nominal global resolution of 2.5 Å (Fig. S5B). Density for the ECD was very poorly resolved in this conformation and could not be resolved with further focused 3D classification, suggesting greater motion of this domain relative to the TMD-G_αs_ portion of the complex (Fig. S5H) as compared to conformer 1. In addition to the ECD, ECL1 was omitted from the model for conformer 2 because of low resolution. Conformer 2 had limited density in the TMD core that could be assigned to the agonist peptide. This density was not sufficient to allow modeling of the ligand (Fig. 3c), suggesting high mobility of the peptide in this receptor conformation.

Comparison of conformers 1 and 2 revealed that the orthosteric pocket of conformer 2 is more open than that of conformer 1 (Fig. 4d, 4e). This structural difference arises from outward motion of the top of TM6 and TM7 in conformer 2 and a more profound kink in the TM6 helix (~100° vs. 71° for conformer 1 vs. 2), which together lead to a ~16 Å outward shift of ECL3 in conformer 2 relative to conformer 1 (Fig. 4c-4e). The position and local conformation of ECL3 in conformer 2 are more similar to ECL3 in the structures reported for the GLP-1R bound to the small molecules TT-OAD2^70^ and CHU-128^12^, which do not contact this loop, than to ECL3 in the structure for the GLP-1R bound to GLP-1^12,40^ or conformer 1 bound with Ex4-D-Ala (Fig. S9).

The weak ligand density in the orthosteric pocket of conformer 2 occurs in a distinct location relative to the ligand bound to conformer 1 (Fig. 4c). Ligand density associated with conformer 2 partially overlaps with the Phe6 and Leu10 side chains of Ex4-D-Ala in conformer 1 but does not appear to extend as deep into the TM core (Fig. 4c). The largest section of continuous density in the orthosteric site of conformer 2 is close to receptor residues Y152^1.47^ and L141^1.36^, which suggests that these residues act as a hydrophobic anchor for the ligand. Y152^1.47^ adopts a distinct rotamer in conformer 2 compared to conformer 1. The ligand density observed in conformer 2 is sterically incompatible with the location of TM1 in conformer 1, which presumably explains why TM1 of conformer 2 is shifted away from the TMD core relative to TM1 in conformer 1 (Fig. 4c, 4e). Weak, transient interactions of Ex4-D-Ala with the receptor core as observed in conformer 2 likely contribute to this state’s high ECD mobility.

The backbone of TM5 is similar in both conformers, but the R310^5.40^ side chain adopts different rotamers. R310^5.40^ is key for receptor activation,^71,72^ and its side chain projects into the orthosteric binding pocket of the TMD in conformer 2. Conversely, the R310^5.40^ side chain projects upwards towards the ECD in conformer 1 (Fig. 4c). Overlaying the two conformers shows that the position of the R310^5.40^ side chain guanidinium group in conformer 2 clashes with the agonist N-terminus in conformer 1 (Fig. 4c). Thus, unfavorable electrostatic and steric interactions would make it impossible for conformer 2 to accommodate the positioning and conformation of the agonist that is observed in conformer 1. Beyond the orthosteric site in the TMD, the agonist, and the ECD, the conformations of the two conformers are largely similar.

The data from 3D classification and the varying local resolution in each of the classes are suggestive of greater conformational dynamics of the GLP-1R when bound to the Ex4-D-Ala peptide relative to that seen with previously solved active, peptide-bound, GLP-1R complexes.^12^ To gain further insight into the dynamics of GLP-1R bound to Ex4-D-Ala, we performed 3D variability analysis in cryoSPARC.^73^ This analysis resolves modes of global motion as principal components, with output of the three major principal components. cryoSPARC analysis identified the transition between conformer 1 and conformer 2 as the dominant principal component within the dataset (Video S1-2). The conformer 2-like state as determined by cryoSPARC again showed unresolved density for the ECD and within the orthosteric pocket. We note that a conformer 2-like state was not previously observed for the GLP-1-bound receptor complex using the same method for 3D variability analysis.^12^ 3D variability analysis also revealed a receptor rocking motion atop the G protein as the major conformational variance in the secondary and tertiary principal components (Video S1-4). Similar dynamic motions of the receptor relative to the heterotrimeric G-protein have recently been detected by cryo-EM for other receptors.^74,75^

## Discussion

Because replacing Gly with D-Ala in a right-handed α-helix involves an energetic penalty of up to 0.5 kcal/mol,^41^ it seems surprising that the major structure we observe via cryo-EM shows the D-Ala residue of Ex4-D-Ala incorporated into the α-helix. It is possible that the D-Ala methyl side chain makes energetically favorable contacts with the receptor that compensate for α-helix destabilization, enabling adoption of the extended N-terminal helix in the G protein-stabilized active state. Nonetheless, the observation that the D-Ala variant binds with ~8-fold lower affinity to the GLP-1R relative to exendin-4 itself suggests that agonist activity is not determined solely by the stability of conformer 1.

We hypothesize that conformer 1 is required for G protein activation, and that adoption of this receptor conformation is favored under conditions used to form a stable complex that can be imaged (inclusion of dominant negative G protein and nanobody 35, apyrase treatment). However, high agonist efficacy might result not only from a propensity to stabilize conformer 1, but also from an ability to promote G protein turnover, which could be hindered if conformer 1 were too long-lived. Dynamics of TMD engagement and release could impact the number of cycles of G protein activation that result from a single agonist-binding event. In the case of Ex4-D-Ala, if the agonist can partially disengage from the TM core but retain other receptor contacts, then the receptor could release the activated G protein and be ready to activate a newly recruited G protein. FRET studies of ligand binding and receptor conformation support the existence of partially and fully engaged ligand-bound states for the PTHR1.^76^ Conformer 2 might represent a partially engaged state, which would presumably occur on the energy surface of the agonist-receptor complex at a position between the completely dissociated and fully bound peptide states (Fig. 5e).

**Fig. 5.**
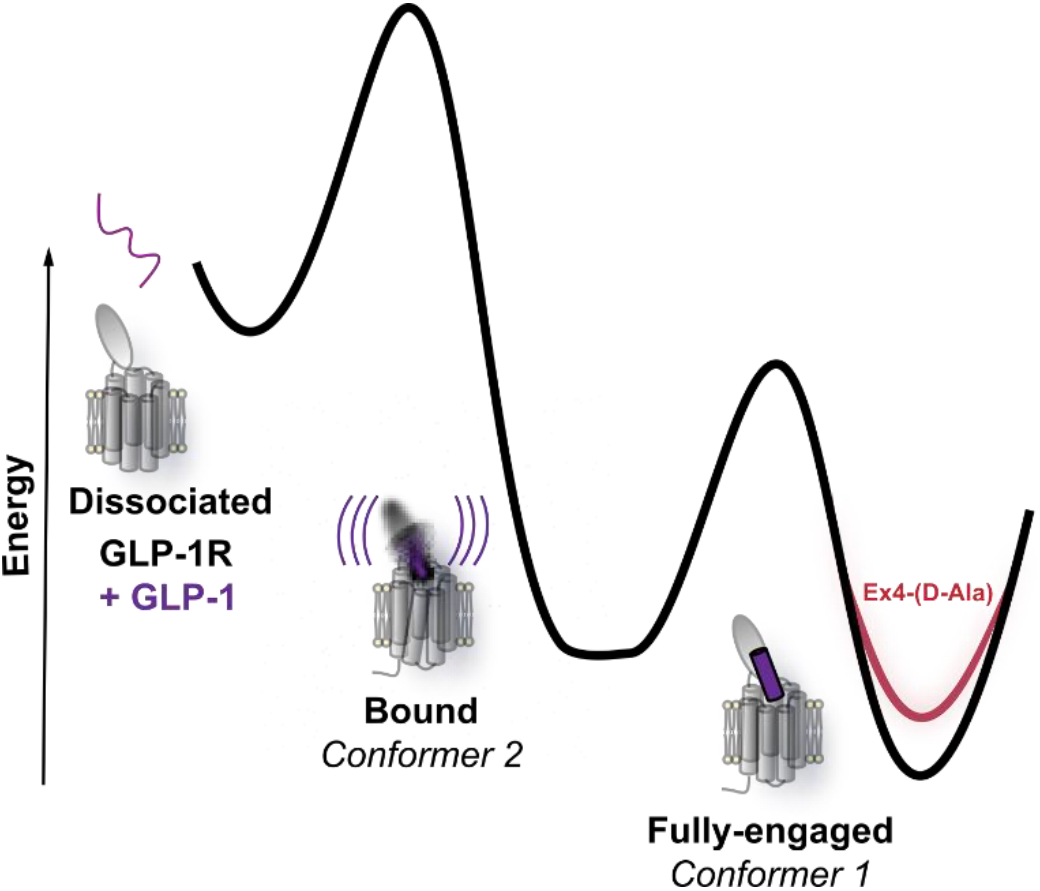
Proposed, simplified energy landscape for the interaction of the GLP-1R and peptide agonists. Dissociated agonist (colored purple) and the GLP-1R are represented as a high energy state at the left. The GLP-1R with fully engaged GLP-1 is represented as the deep energy well at the right. Because the Gly10→D-Ala analogue of exendin-4 binds to the GLP-1R with lower affinity than does GLP-1, the energy well for the fully engaged state in this case (shown in red) is higher than for GLP-1. The central energy well reflects a bound state of intermediate stability that features significant internal motion and corresponds to conformer 2.

The plausibility of this hypothesis is indirectly supported by the observation that salmon calcitonin, a high-affinity agonist, displays slower G protein release kinetics, and thus lower efficacy, than human calcitonin, a lower-affinity agonist.^62^ These two agonists favor different G protein conformations at the CTR, as indicated by the lower BRET signal in the G protein conformational change assay for human relative to salmon calcitonin.^62^ Similarly, we observed that Ex4-D-Ala induced a lower maximal BRET signal relative to GLP-1 in a comparable assay (Fig. 2d).

The stability of a partially engaged state, the stability of the fully bound state (which is competent for G protein activation) and the height of the energy barrier separating these two states could all be affected by changes at Gly10 of GLP-1 or Gly4 of exendin-4 (Fig. 5), and changes in these factors might explain the variations in behavior observed among the set of peptides studied here. We propose that the poor efficacy of the analogues containing (*S,S*)-ACPC arises because this residue stabilizes the helical conformation near the N-terminus, relative to the native Gly, and thereby raises the energy barrier between the partially engaged and fully engaged states. Hindered exchange between the fully and partially engaged states might prevent the activation of multiple G proteins after a single agonist-receptor association event. For the analogues containing D-Ala, on the other hand, disfavoring helicity near the N-terminus might lower the barrier for interconversion between the partially and fully engaged states, and thereby enhance the likelihood that multiple G protein activation cycles would be triggered by a single agonist-receptor association. In this case, the diminished affinities of the D-Ala analogues relative to the natural agonists could be compensated by an increase in average number of G proteins activated to cause the observed similarity in receptor activation efficacies of the D-Ala analogues relative to GLP-1 and exendin-4.

We cannot rule out the possibility that either of our cryo-EM-derived conformers, alone, represents the signal-transducing form of the agonist-receptor complex, and that the other conformer lacks functional significance. The alternative hypothesis offered above, however, is consistent with previous studies that support a role for ligand mobility in activation of other GPCRs. Dynorphin, a short opioid peptide, retains disorder when bound to the kappa opioid receptor,^77^ and mobility of neurotensin residue Tyr-11 is required for activation of the cognate receptor.^78^ Receptor activity-modifying proteins (RAMPs) also alter the conformational dynamics of the adrenomedullin receptors to effect changes in receptor phenotype.^74^ Our findings are distinct from these precedents, however, in suggesting that at least two distinct states of an agonist-receptor complex may play important and complementary roles in the signal transduction mechanism.

Collectively, the data reported here suggest that interconversion among distinct agonist-receptor conformations is critical to the efficacy of signal-transduction via the GLP-1R. This conclusion is consistent with emerging evidence that conformational mobility in agonist and receptor can be functionally important in signal-transducing states of other GPCR-peptide complexes.^25,77,78^ The mode of agonist mobility highlighted in this work may be evolutionarily conserved among peptide agonists of related Class B1 GPCRs; Gly at the fourth position from the N-terminus is found in glucagon, GLP-2 and several other hormones.^33^ Other sites of essential mobility may be present in more distantly related hormones. For example, both parathyroid hormone (PTH) and parathyroid hormone related protein have Gly at position 12, and early work suggested that PTH activity is retained when Gly12 is replaced by either D-Ala or L-Ala.^79^ Understanding the role of structural dynamics in the propagation of molecular information across the cell membrane is important in terms of elucidating GPCR function and developing improved therapeutic agents. A dynamics-based approach to drug design would represent a departure from traditional approaches, which focus on promoting a specific conformation rather than retaining or enhancing particular modes of conformational mobility that might contribute to efficacy by mechanisms other than high-affinity binding. A deeper understanding of the conformational possibilities available to GPCRs bound to flexible agonists, and of relationships among conformational states and signal transduction, will enhance prospects for elucidating signal-propagating mechanisms at the molecular level and optimizing therapeutic performance.

## Author Contributions

B.P.C. designed the project, synthesized peptides, and expressed and purified protein complex. B.P.C., P.Z., and T.T.T. conducted in vitro assays. B.P.C., S.J.P., and M.J.B. processed the cryo-EM data, built the model, and performed refinement. S.J.P. and M.J.B. performed the multivariate analysis, assisted in data interpretation, and assisted in figure preparation. R.D. prepared the cryo-EM sample and collected EM data. P.M.S., D.W., and S.H.G. supervised the project. B.P.C., P.M.S., D.W., and S.H.G. interpreted data, generated figures, and wrote the manuscript. All authors reviewed and edited the manuscript.

## Acknowledgements

This work was supported by the National Institutes of Health (R01 GM056414, to S.H.G.). B.P.C. was supported in part by a graduate fellowship from the NSF (DGE-1747503). B.P.C. was supported in part by a Biotechnology Training Grant from NIGMS (T32 GM008349). R.D. was supported by Takeda Science Foundation 2019 Medical Research Grant and Japan Science and Technology Agency PRESTO (18069571). P.M.S. and D.W. were supported by an ARC Centre Grant (IC200100052). P.M.S. was supported by the National Health and Medical Research Council of Australia (NHMRC) Program Grant (1150083), and Senior Principal Research Fellowship (1154434). D.W. was supported by NHMRC Project Grants (1126857 and 1184726), and a NHMRC Senior Research Fellowship (1155302).

