## Supplementary Text and Figures for "Structural and Functional Diversity among Agonist-Bound States of the GLP-1 Receptor"

**Figure S1**

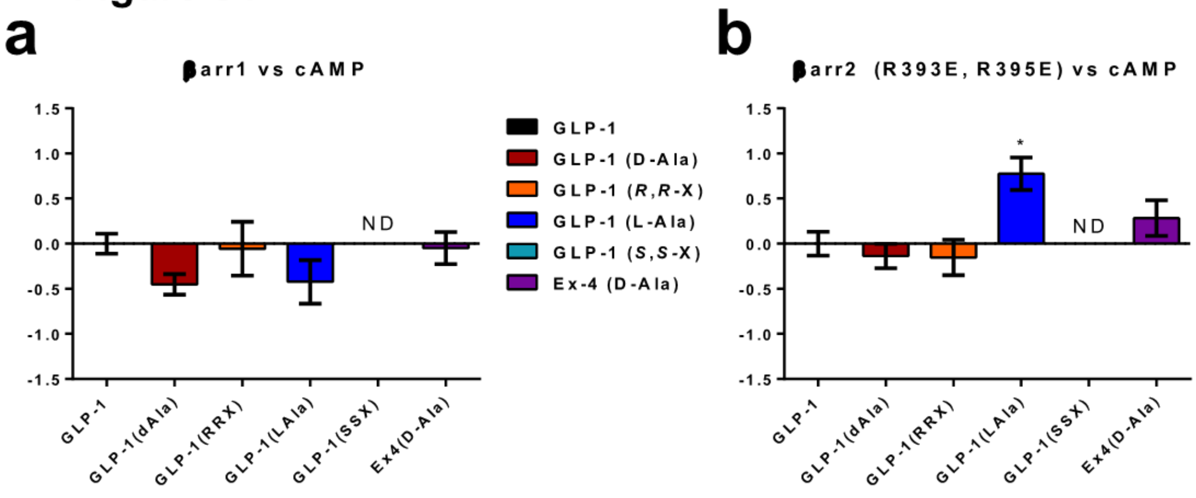

Fig. S1 | Bias factors calculated from normalization of the Log(tau/$K_A$) ratios, determined by the Black-Leff operation model, to the reference agonist GLP-1 and reference pathway, cAMP. The native agonist, GLP-1, is assigned a bias factor of 0 by definition. Positive values indicate bias towards arrestin recruitment, and negative values indicate bias towards cAMP production. Error bars represent standard deviation. * indicates $P < 0.05$ ($P = 0.026$) compared to GLP-1 by one-way analysis of variance (ANOVA) with Dunnett's post-test. No data (ND) signifies that a bias factor could not be computed (**A**) β-arrestin-1 vs cAMP bias factors (**B**) β-arrestin-2 (R393E, R395E) vs cAMP bias factors.

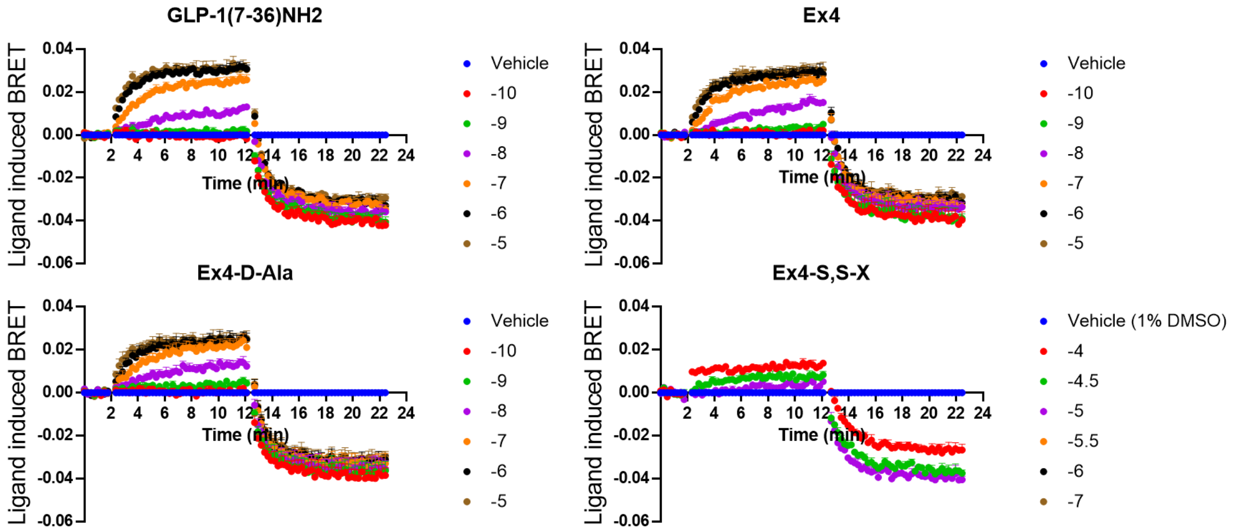

Fig. S2 | G protein conformation assay time-course. The ligand induced BRET is baseline-subtracted. Agonist was added at 2 min and GTP (30 µM) was added after the 12 min timepoint. Different colors indicate different concentrations (log[peptide (M)]), as indicated by the key on the right side of each plot. Data points represent the mean of three independent experiments. Error bars represent standard error of the mean.

#### Figure S3

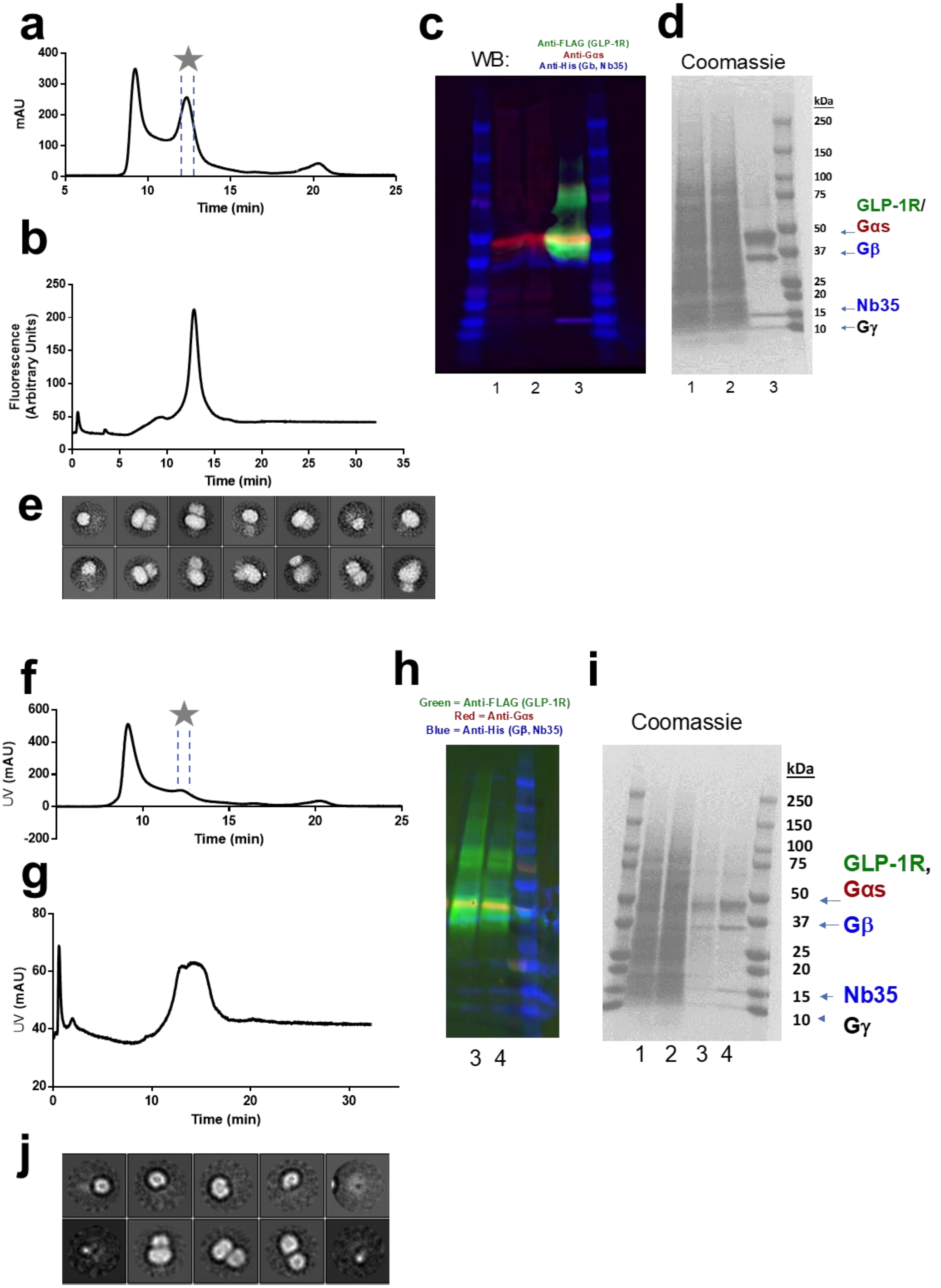
a
mAU
Time (min)
b
Fluorescence (Arbitrary Units)
Time (min)
c
WB:
Anti-FLAG (GLP-1R)
Anti-Gαs
Anti-His (Gb, Nb35)
1 2 3
d
Coomassie
kDa
250
150
100
75
50
37
25
20
15
10
GLP-1R/
Gαs
Gβ
Nb35
Gγ
1 2 3
e
f
UV (mAU)
Time (min)
g
UV (mAU)
Time (min)
h
Green = Anti-FLAG (GLP-1R)
Red = Anti-Gαs
Blue = Anti-His (Gβ, Nb35)
3 4
i
Coomassie
kDa
250
150
100
75
50
37
25
20
15
10
GLP-1R,
Gαs
Gβ
Nb35
Gγ
1 2 3 4
j

Fig. S3 | **a-e**; Purification and characterization of the GLP-1R/Ex4-D-Ala/DNGαs/Gβ1/Gγ2/Nb35 complex. **a**, Size-Exclusion chromatogram of crude anti-FLAG elution. **b**, Fluorescence-detected size-exclusion chromatogram of purified complex. **c**, Western blot. The channel corresponding to anti-His6 antibody is depicted in blue. The channel corresponding to anti-FLAG antibody channel is depicted in green. The channel corresponding to anti-Gas antibody is depicted in red. **d**, Coomassie-stained, reducing SDS-PAGE gel. The lanes labeled 1, 2, and 3 correspond to LMNG/CHS solubilized fraction, anti-FLAG column flow-through, and purified complex, respectively for both **c** and **d**. **e**, Representative 2D classes from negative-stain, single-particle electron microscopy of the GLP-1R/Ex4-(D-Ala)/DNGαs/Gβ1/Gγ2/Nb35 complex. **f-j**; Purification and characterization of the GLP-1R/Ex4-S,S-X/DNGαs/Gβ1/Gγ2/Nb35 complex. **f**, Size-Exclusion chromatogram of crude anti-FLAG elution. **g**, Fluorescence-detected size-exclusion chromatogram of purified complex. **h**, Western blot. The channel corresponding to anti-His6 antibody is depicted in blue. The channel corresponding to anti-FLAG antibody channel is depicted in green. The channel corresponding to anti-Gαs antibody is depicted in red. **i**, Coomassie-stained, reducing SDS-PAGE gel. The lanes labeled 1, 2, 3, and 4 correspond to LMNG/CHS solubilized fraction, anti-FLAG column flow-through, anti-FLAG elution, and purified complex, respectively for both **h**, and **i**. **j**, Representative 2D classes from negative-stain, single-particle electron microscopy of the GLP-1R/Ex4-(S,S-X)/DNGαs/Gβ1/Gγ2/Nb35 complex.

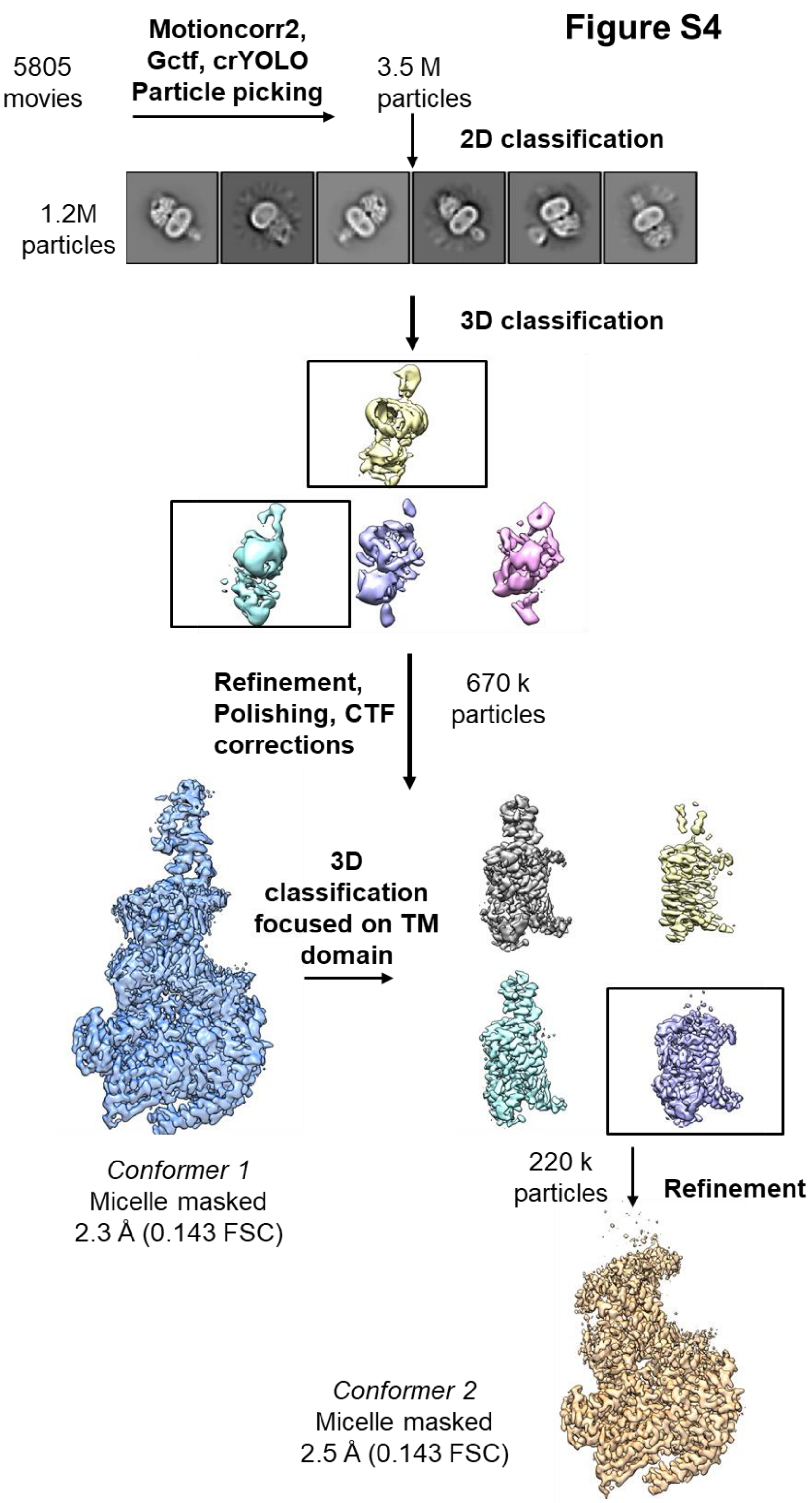

Fig. S4 | An overview of the cryo-EM data processing pipeline for the Ex4-D-Ala/GLP-1R/Gs complex.

**Figure S5**

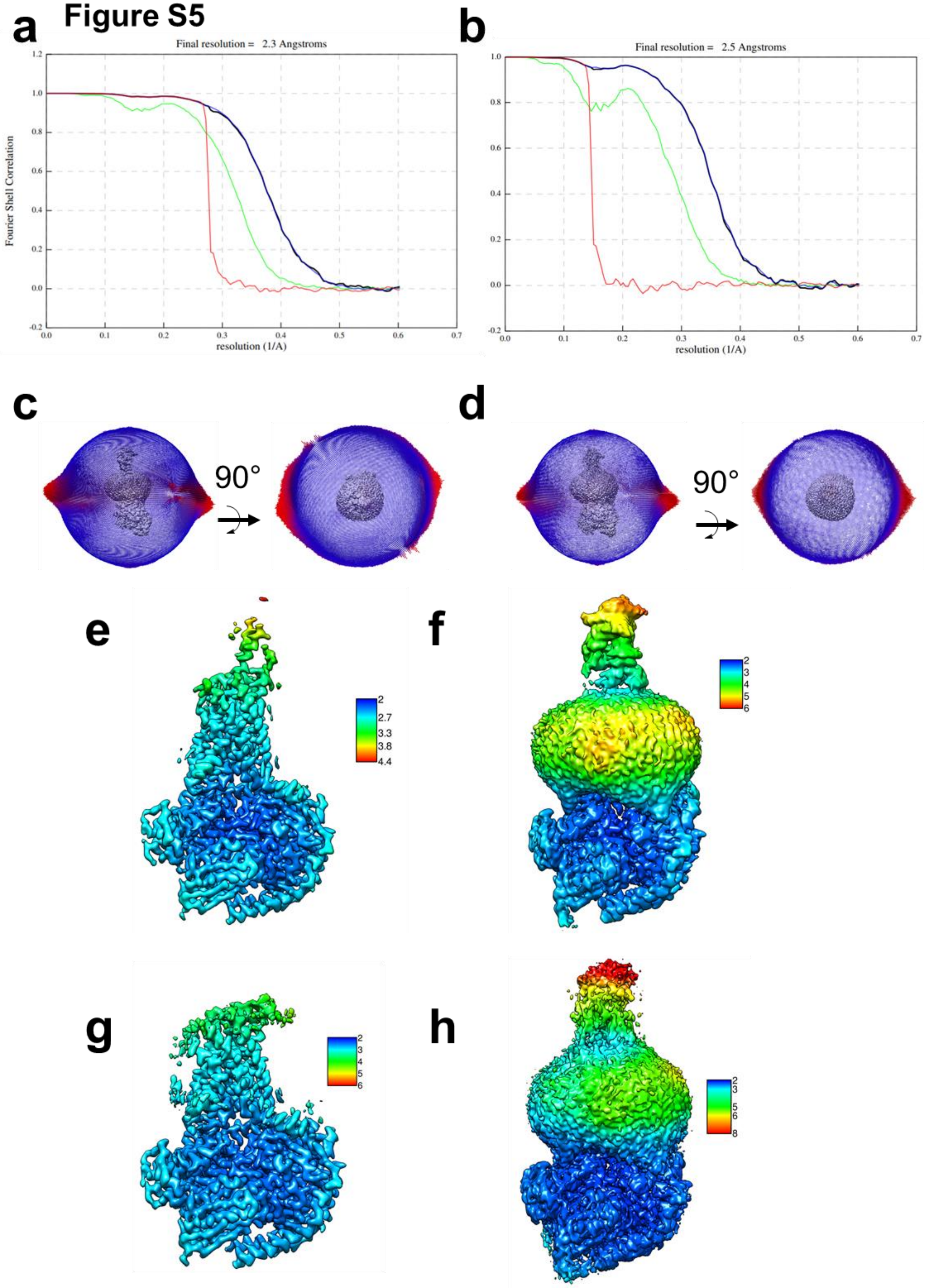

a
Final resolution = 2.3 Angstroms
Fourier Shell Correlation
resolution (1/A)
b
Final resolution = 2.5 Angstroms
resolution (1/A)
c
90°
d
90°
e
2
2.7
3.3
3.8
4.4
f
2
3
4
5
6
g
2
3
4
5
6
h
2
3
5
6
8

Fig. S5 | **a-b**, Gold-standard Fourier Shell Correlation (FSC) curves for conformers 1 and 2 showing overall nominal resolutions of 2.3 Å and 2.5 Å, respectively. Black, green, blue and red curves indicate corrected, unmasked, masked, and phase-randomized maps, respectively. **c-d**, Euler angle distribution histograms of the particles used in reconstructions. **c,** Conformer 1 **d,** Conformer 2 **e-h**, Local resolution estimates (computed with RELION 3.1 Beta) are shown as colored heatmaps. **e**, Conformer 1 at low threshold. **f**, Conformer 1 at high threshold. **g**, Conformer 2 at low threshold. **h**, Conformer 2 at high threshold.

#### Figure S6

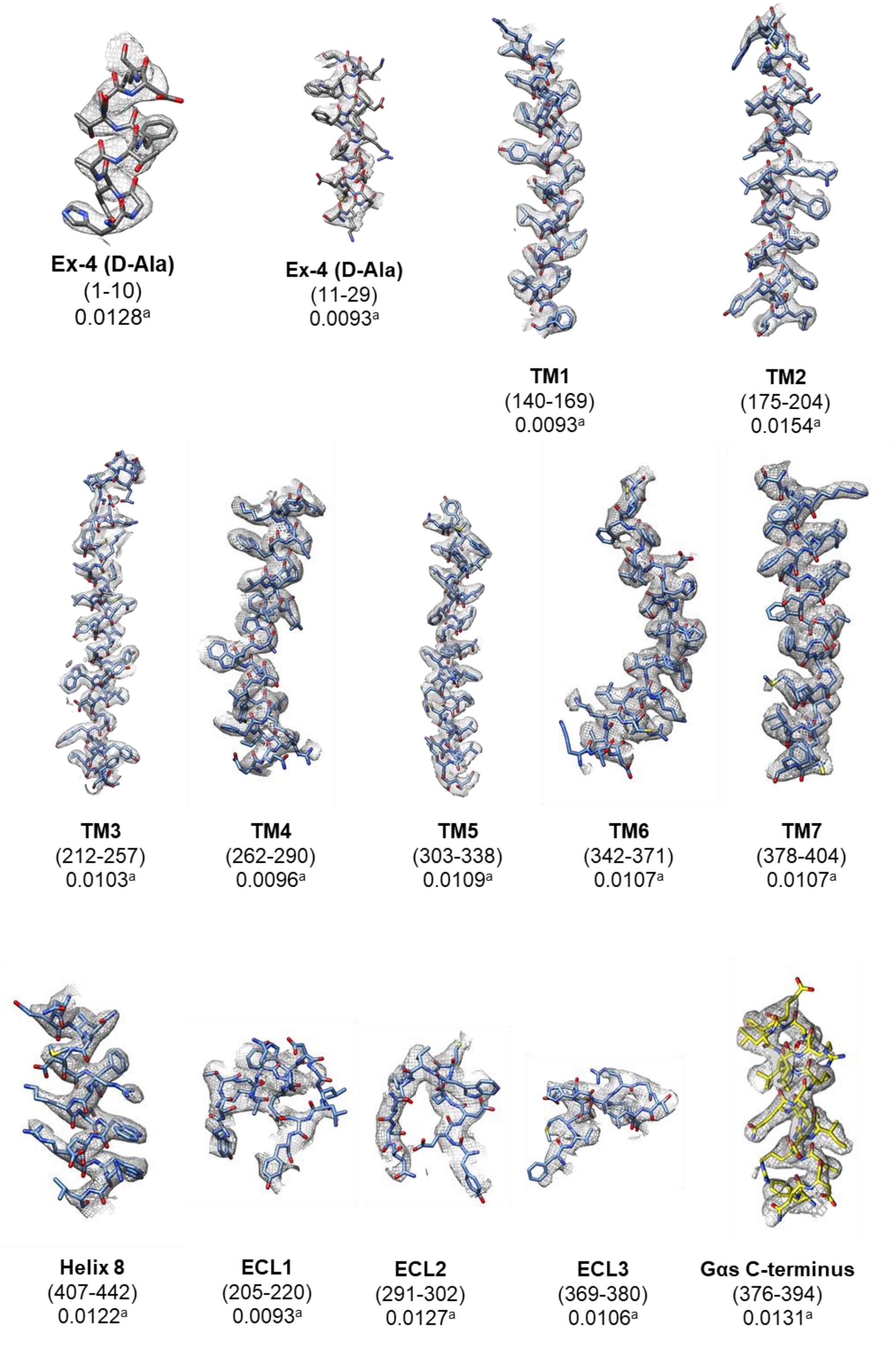

Fig. S6 | The electron microscopy density map and the model are shown for selected features of Conformer 1. Residue numbers are show in parentheses. [a] indicates the map threshold.

#### Figure S7

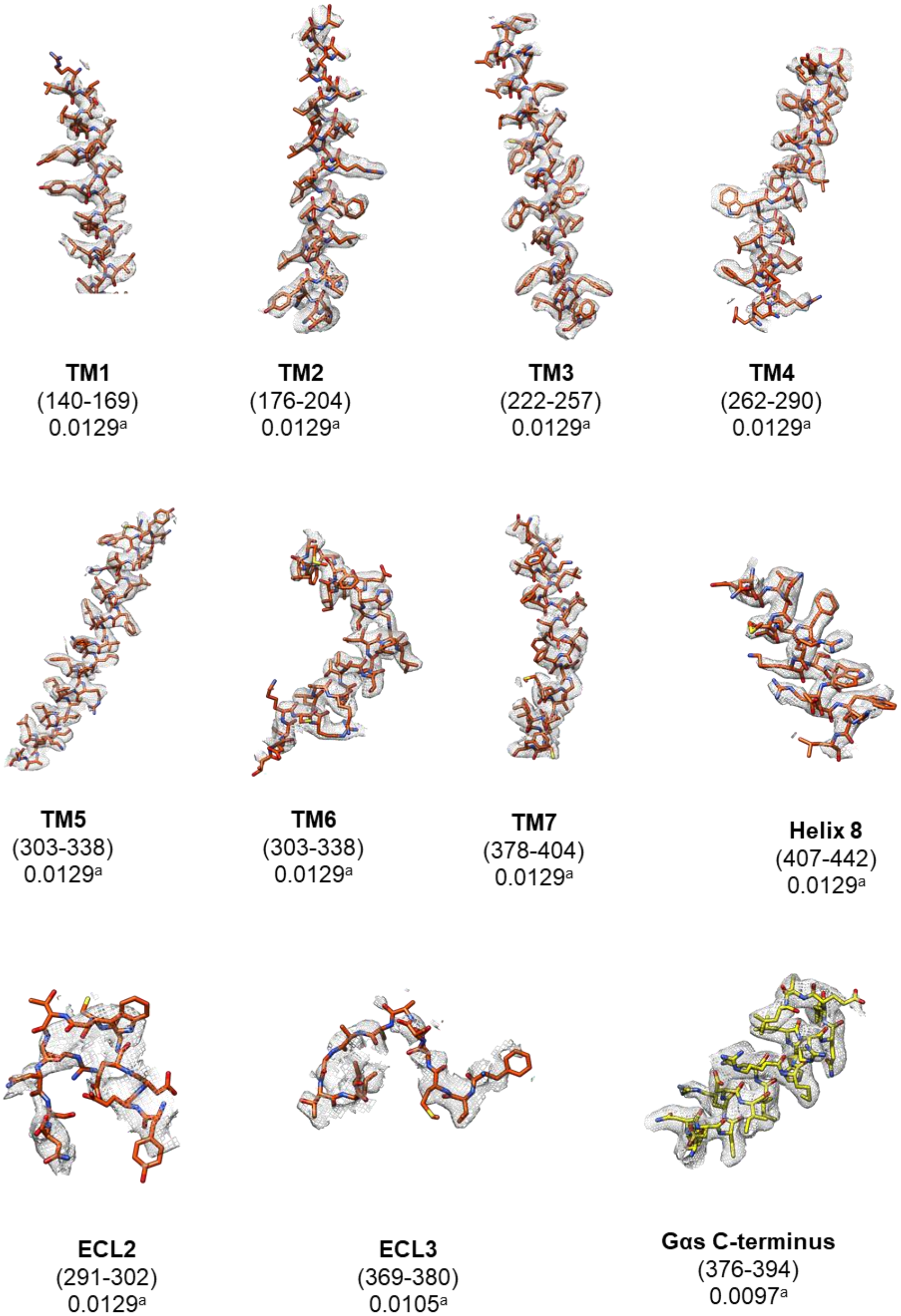

Fig. S7 | The electron microscopy density map and the model are shown for selected features of the conformer 2 complex. Residue numbers are show in parentheses. [a] indicates the map threshold.

**a** **Figure S8**

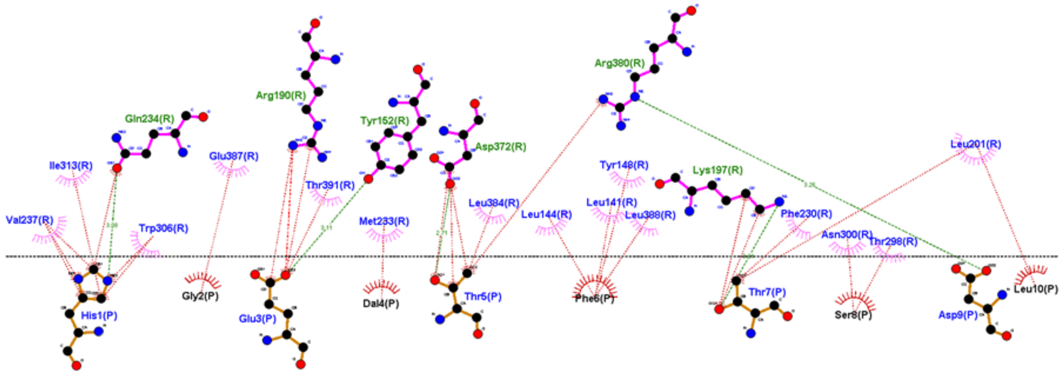

**b**

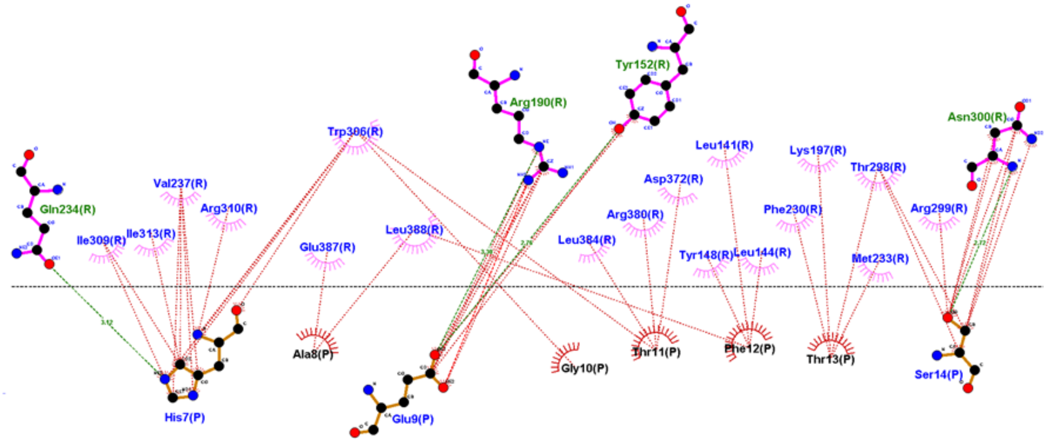

Fig. S8 | A Ligplot+ v2.2 diagram of the N-terminal interacting residues of Ex-4-D-Ala bound to GLP-1R as observed in the conformer 1 model and GLP-1 bound to GLP-1R (A and B, respectively). Residues of the agonist are indicated by the chain (P) denotation and GLP-1R residues are indicated by the (R) denotation. Hydrophobic interactions are shown with dotted red lines and hydrogen bonding is indicated with dotted green lines. Hydrogen bonds are shown with a maximum distance of 3.35 Å and other non-bonded contacts are shown with 3.90 Å.

Figure S9

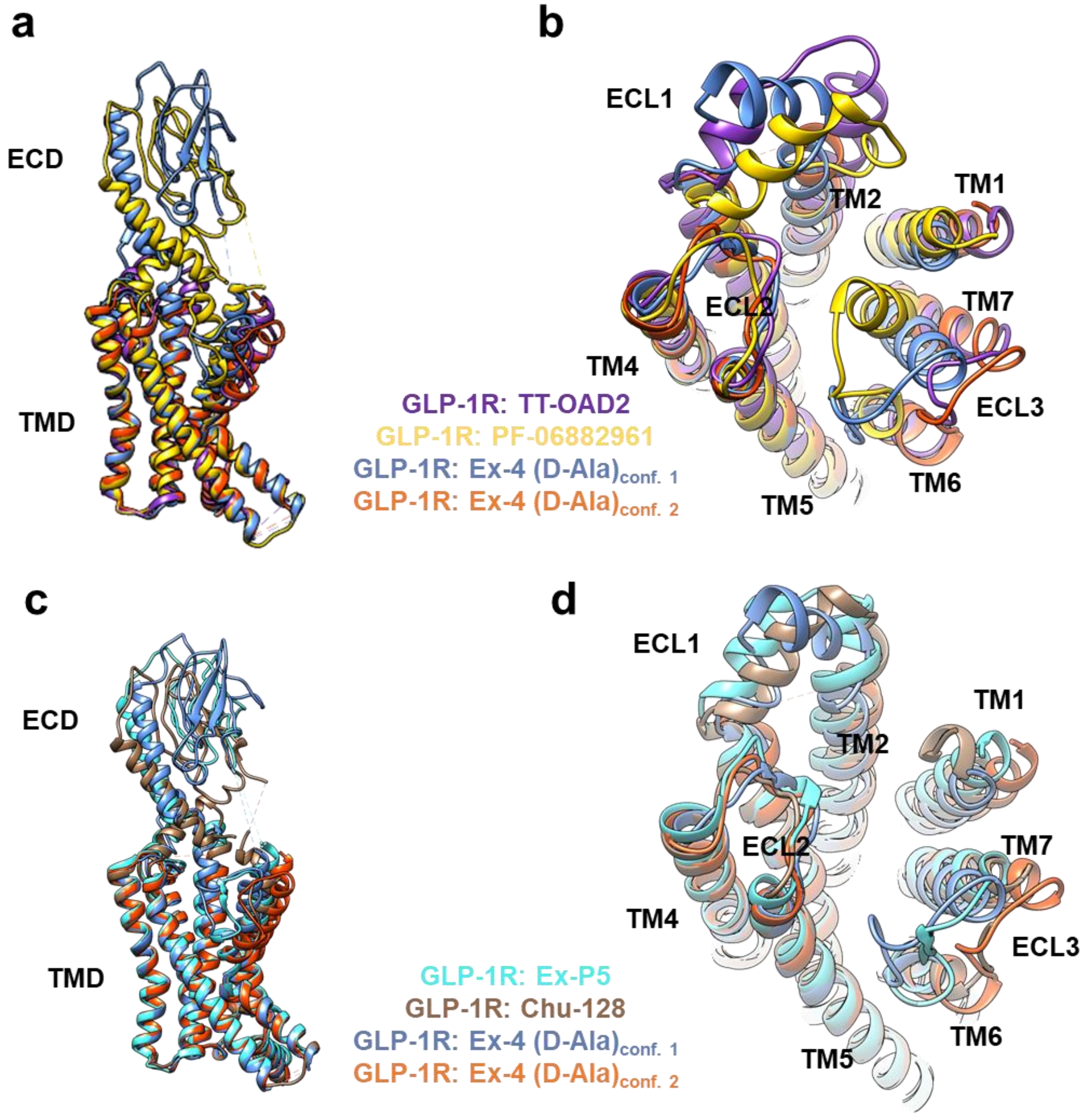

Fig. S9 | A comparison of GLP-1R structures with ligand removed (and ECD removed in top views) for clarity. (A) Side view and (B) top view of GLP-1R bound to TT-OAD2, PF-06882961, and Ex-4-(D-Ala)'s two conformers shown in purple, yellow, blue and orange, respectively. (C) Side view and (D) top view of GLP-1R bound to Ex-P5, Chu-128 and Ex-4-(D-Ala)'s two conformers shown in cyan, brown, blue, and orange, respectively.

### Table S1

| | cAMP Production Inhibition | | |
|---|---|---|---|
| | $pIC_{50}$ | $IC_{50}$(nM) | $IC_{50}$rel. |
| **Ex-(9-39)** | 7.20 ± 0.06 | 63 | 1 |
| **GLP-1-S,S-X** | 7.38 ± 0.05 | 42 (16%[a]) | 0.67 |
| **Ex4-S,S-X** | 7.85 ± 0.04 | 14 (1.2%[a]) | 0.22 |

Table S1 | Fit parameters for concentration-response data in main text Figure 2a. $IC_{50}$ rel. indicates the potency relative to Exendin-(9-39) by the quotient (Peptide $IC_{50}$) / (Ex-9-39 $IC_{50}$) [a] indicates the maximal cAMP response in the absence of GLP-1 addition. Uncertainties are expressed as standard error of the mean. N = 3.

### Table S2

| | βArrestin-1 Recruitment | | | | | βArrestin-2 Recruitment | | | | |
|---|---|---|---|---|---|---|---|---|---|---|
| | $pEC_{50}$ | $EC_{50}$ (nM) | % Max | $EC_{50\ rel.}$ | Bias Factor | $pEC_{50}$ | $EC_{50}$ (nM) | % Max | $EC_{50\ rel.}$ | Bias Factor |
| **GLP-1** | 8.36 ± 0.1 | 4.4 | 100 ± 6 | 1 | 0 ± 0.2 | 8.31 ± 0.2 | 4.9 | 99 ± 7 | 1 | 0 ± 0.2 |
| **GLP-1-D-Ala** | 7.85 ± 0.1 | 14 | 86 ± 6 | 3.2 | -0.45 ± 0.2 | 8.14 ± 0.1 | 7.2 | 81 ± 5 | 1.5 | -0.1 ± 0.2 |
| **GLP-R,R-X** | 7.43 ± 0.3 | 38 | 18 ± 3 | 8.5 | -0.06 ± 0.5 | 6.84 ± 0.4 | 140 | 49 ± 13 | 29 | -0.2 ± 0.3 |
| **GLP-1-L-Ala** | 7.17 ± 0.3 | 67 | 24 ± 4 | 30 | -0.4 ± 0.4 | 8.05 ± 0.2 | 8.9 | 46 ± 4 | 1.8 | 0.8 ± 0.3* |
| **GLP-1-S,S-X** | - | - | - | - | - | - | - | - | - | - |
| **Ex4-D-Ala** | 8.10 ± 0.2 | 7.9 | 58 ± 4 | 1.8 | -0.05 ± 0.3 | 8.31 ± 0.2 | 4.9 | 68 ± 7 | 1 | 0.3 ± 0.3 |

Table S2 | $EC_{50}$ values, maximal responses, and $IC_{50}$ values from 3-parameter sigmoidal fits for concentration-response data in figures 2**b** and 2**c**. $EC_{50}$ rel. indicates β-arrestin recruitment potency relative to GLP-1 by the quotient (Peptide $EC_{50}$) / (GLP-1 $EC_{50}$). Uncertainties are expressed as standard error of the mean. N = 3. Bias factors as determined by the Black-Leff operation model. The native agonist, GLP-1, is assigned a bias factor of 0 by definition. Positive values indicate bias towards arrestin recruitment, and negative values indicate bias towards cAMP production. * indicates $P < 0.05$ ($P = 0.026$) compared to GLP-1 by one-way analysis of variance (ANOVA) with Dunnett's post-test.

### Table S3

| | G-protein conformational change | | | | |
|---|---|---|---|---|---|
| | $pEC_{50}$ | $EC_{50}$ (nM) | % Max | $EC_{50\,rel.}$ | Rate at 100 nM ($min^{-1}$) |
| **GLP-1** | 7.84 ± 0.1 | 14 | 100 ± 4 | 1 | 0.48 ± 0.03 |
| **Ex4** | 8.1 ± 0.1 | 8.5 | 95 ± 4 | 0.6 | 0.42 ± 0.03 |
| **Ex4-D-Ala** | 8.0 ± 0.2 | 9.4 | 80 ± 5 | 0.7 | 0.40 ± 0.04 |
| **Ex4-*S*,*S*-X** | <5 | >10,000 | 45 ± 2[a] | >1000 | ND |

Table S3 | Fit parameters for concentration-response data in main text figure 3. $EC_{50}$ rel. indicates G-protein potency relative to GLP-1 by the quotient (Peptide $EC_{50}$) / (GLP-1 $EC_{50}$). Rates were determined by fitting kinetic data at 100 nM ligand concentration to one-phase association. Uncertainties are expressed as standard error. ND indicates no data [a] Indicates the response at 100 µM ligand concentration. Ex4-*S*,*S*-X did not reach saturation.

### Table S4

| **Imaging** | *Ex-4-D-Ala/GLP-1R/Gs* | |
|---|---|---|
| Magnification | 105,000 | |
| Voltage | 300 kV | |
| Spot Size | 5 | |
| Electron exposure (e-/Å2 ) | 49.8 | |
| Exposure time (s) | 5.011 | |
| Movie frames | 71 | |
| Pixel size (Å) | 0.83 | |
| K3 CDS mode | Yes | |
| **Processing** | *Conformer 1* | *Conformer2* |
| Symmetry imposed | C1 | C1 |
| Final particle imaged (no.) | 671,599 | 221,955 |
| Resolution (Å) | 2.32 | 2.51 |
| FSC threshold | 0.143 | 0.143 |
| **Refinement** | | |
| Initial model used (PDB code) | 6B3J | 6B3J |
| Map sharpening B factor (Å2 ) | -10 | -10 |
| **Model composition** | | |
| Chains | 6 | 5 |
| Non-Hydrogen Atoms | 9154 | 8097 |
| Protein residues | 1170 | 1023 |
| Ligand | 1 | 0 |
| **RMSDs** | | |
| Bond length (Å) | 0.007 | 0.008 |
| Bond angles (°) | 0.973 | 0.826 |
| **Validation** | | |
| MolProbity score | 1.32 | 1.42 |
| Clashscore | 5.89 | 6.15 |
| Rotamer outliers (%) | 0.63 | 0.23 |
| **Ramachandran plot** | | |
| Favored (%) | 98.34 | 97.61 |
| Allowed (%) | 1.66 | 2.39 |
| Disallowed (%) | 0 | 0 |

Table S4 | Cryo-EM imaging, model composition and validation parameters. The map sharpening B factors were manually applied.

### Materials and Instrumentation:

#### A. Instrumentation:

Automated solid-phase peptide synthesis (SPPS) was performed with a CEM Liberty Blue instrument with an $HT_{12}$ resin-handling module. Manual, microwave assisted SPPS was performed with Torviq polypropylene syringes fitted with a porous polypropylene disk at the bottom using a CEM MARS microwave instrument. Preparative HPLC was performed using either a Shimadzu HPLC system (SCL-10VP system controller, LC-6AD pumps, SIL-10ADVP autosampler, SPD-10VP UV-vis detector, FRC-10A fraction collector) or a Waters HPLC system (Waters 2545 Binary Gradient module, Waters 2707 Autosampler, Waters 2998 Photodiode Array Detector, and Waters Fraction Collector III). The HPLC systems were outfitted with a C18 column (either Agilent ZORBAX 21.2x250 mm, 7 µm or Waters XSelect CSH 19x250 mm, 5 µm). Matrix-assisted laser desorption ionization-time-of-flight (MALDI-TOF) mass spectrometry was performed by pipetting 1 µL of peptide mixture and 1 µL of CHCA (saturated solution in 1:1 acetonitrile:water) onto an appropriate target, allowing the liquid to evaporate completely under ambient conditions, and acquiring spectra on a Bruker microflex™ LRF. Liquid-Chromatography-Mass Spectrometry (LCMS) was performed on a Waters Acquity Arc system outfitted with a Waters XBridge C18 1.2 x 50 mm (3.5 µm) column. UV-Vis measurements of peptides and plasmids were performed with with either a ThermoFisher NanoDrop 2000 or a NanoDrop One C microvolume spectrophotometer. Luminescence and bioluminescence resonance energy transfer (BRET) were measured with a BioTek Synergy 2 plate reader. Size-exclusion chromatography was performed with a Bio-rad system equipped with a GE Superdex 200 Increase 1G/300 GL column, BioLogic Dual flow pump, BioFrac Fraction Collector, and a Shimadzu RF-10AXL Fluorescence detector. Western blots were imaged with an Amersham Typhoon 5 imager. Negative-stain electron microscopy was performed with a ThermoFisher Talos L120C transmission electron microscope, and grids for negative staining were glow-discharged with a Quorum GloQube Plus.

#### B. Instrument Acknowledgements:

The purchase of the Bruker microflex™ LRF in 2015 was funded by the Bender gift to the Department of Chemistry.

#### C. Table of Materials:

| Item/Reagent | Vendor | Item # | Notes |
|---|---|---|---|
| **α-N-Fmoc-amino acids** | Chem Impex | Various | |
| **Fmoc-(*S*,*S*)-ACPC** | Chem Impex | 15073 | |
| **Fmoc-(*R*,*R*)-ACPC** | Chem Impex | 14468 | |
| **NovaPEG Rink Amide** | Milipore-Sigma | 8550470005 | |
| **SPPS Reaction Vessel** | Torviq | SPE-0500 | (5mL) |
| **O-(7-Azabenzotriazol-1-yl)-N,N,N',N'-tetramethyluronium hexafluorophosphate (HATU)** | Chem Impex | 12881 | |
| **N,N'-Diisopropylcarbodiimide (DIC)** | Chem Impex | 00110 | |

| | | | |
|---|---|---|---|
| **Ethyl (hydroxyimino)cyanoacetate (Oxyma)** | Chem Impex | 26426 | |
| **α-Cyano-4-hydroxycinnamic acid (CHCA)** | Sigma-Aldrich | 70990 | |
| **Diisopropylethylamine (DIEA)** | Sigma-Aldrich | 496219 | Biotech. Grade |
| ***N*, *N* Dimethylformamide (DMF)** | Sigma-Aldrich | 319937 | ACS Reagent |
| ***N*, *N* Dimethylformamide (DMF)** | Sigma-Aldrich | 494488 | Biotech. Grade |
| **Piperidine** | Sigma-Aldrich | 104094 | |
| **Piperazine** | Sigma-Aldrich | P45907 | |
| **Trifluoroacetic acid** | Sigma-Aldrich | T6508 | ReagentPlus, 99% |
| **1,2 Ethanedithiol** | Sigma-Aldrich | 2390 | |
| **Thioanisole** | Sigma-Aldrich | T28002 | |
| **D-Luciferin** | GoldBio | LUCK-100 | (Potassium Salt) |
| **Coelenterazine 400 a** | GoldBio | C-320-10 | also known as DeepBlueC™ |
| **H-Coelenterazine** | Nanolight | 301 | |
| **FuGENE® HD** | Promega | E2311 | |
| **Nano-Glo® Luciferase Assay System** | Promega | N1110 | |
| **pCMV6-XL5 [hGLP1-R]** | Origene | SC124060 | |
| **pSectag2 (modified) [GLP-1R-His-Flag]** | - | - | Kind gift from Adam Schwaid (Pfizer)[1] |
| **nLuc-GLP-1R-His-FLAG** | This paper | Available at Addgene (# 124831) | |
| **GLP-1R-Rluc8** | In house | - | See Hager et al.[2] |
| **pCMV6-XL5 [GFP$^2$ß-Arrestin-1]** | In house | - | See Hager et al.[2] |
| **GFP$^2$ß-Arrestin-2(R393E, R395E)** | In house | - | See Hager et al.[2] |
| **Custom DNA oligonucleotides** | Integrated DNA Technologies | - | 25 nmole DNA oligo, standard desalting |
| **GRK5** | - | - | Kind Gift from Rasmus Jorgensen and Jakob Lerche Hansen (Novo Nordisk) |
| **HEK293FT Cells** | ThermoFisher | R70007 | |
| **HEK293-GS22 Cells** | Promega | E1261 | Kind Gift from Thomas Gardella |
| **DMEM** | Gibco (ThermoFisher) | 11965092 | +High glucose, L-Gln, Phenol Red |
| **Opti-MEM** | Gibco (ThermoFisher) | 31985070 | With L-Gln |
| **McCoy's 5A Medium Modified** | Gibco (ThermoFisher) | 12330031 | +High glucose, L-Gln, Bacto-peptone, Phenol Red, Hepes |
| **DPBS** | Sigma-Aldrich | D8537 | |
| **DPBS (+Glucose)** | Gibco (ThermoFisher) | 14287080 | +calcium, magnesium, glucose, pyruvate |
| **100X L-Glutamine** | Gibco (ThermoFisher) | 25030081 | |
| **100X Sodium Pyruvate** | Gibco (ThermoFisher) | 11360070 | |
| **100X MEM NEAA** | Gibco (ThermoFisher) | 11140050 | |

| | | | |
|---|---|---|---|
| **Hyclone™ 100X Penicillin-Streptomycin** | GE Healthcare Life Sciences | SV30010 | |
| **Fetal Bovine Serum** | Gibco | 10082147 | Heat Inactivated |
| **10 cm, tissue-treated dishes** | Corning | 353003 | |
| **75 cm² vented, tissue-treated flasks** | Corning | 430641U | |
| **0.05% Trypsin-EDTA** | Gibco (ThermoFisher) | 25300054 | With Phenol Red |
| **96-well plates (white bottom)** | Corning | 3917 | TC-treated, white, sterile |
| **Phusion High-Fidelity PCR kit** | Thermo | F530S | |
| **DNA Cleanup kit** | Zymo | D4013 | |
| **Gibson Assembly® Cloning Kit** | New England Biolabs | E5510S | |
| **EndoFree Plasmid Maxi Kit** | Qiagen | 12362 | |
| **Human GLP-1(7-36)-$NH_2$** | Prospec | HOR-284 | |
| **Exendin-4-$NH_2$** | Anaspec | AS-24463 | |
| **Exendin (9-39)-$NH_2$** | Genscript | RP10872 | |
| **ESF 921™ Insect Cell Culture Medium** | Expression Systems | 96-001-01 | |
| **4–15% Mini-PROTEAN® TGX™ Precast Protein Gels** | BioRad | 4561086 | |
| **Anti-FLAG M1 Affinity Resin** | Millipore Sigma | A4596 | |
| **EGTA** | Combi-Blocks | QB-6401 | |
| **Anti-FLAG M1 antibody** | Millipore Sigma | F3040 | Mouse monoclonal |
| **Anti-GNAS antibody** | US Biological Life Sciences | 036131 | Rabbit polyclonal |
| **Anti-$His_5$ antibody** | Qiagen | 34660 | Mouse monoclonal |
| **IRDye® 680RD Goat-anti-Mouse Antibody** | Li-Cor | 926-68070 | |
| **IRDye® 800CW Goat-anti-Rabbit Antibody** | Li-Cor | 926-32211 | |
| **Precision Plus Protein™ Dual Color Standards** | BioRad | 1610374 | |
| **Apyrase** | New England Biolabs | M0398L | |
| **Lauryl Maltose Neopentyl Glycol (LMNG)** | Anatrace | NG310 | |
| **Cholesteryl Hemisuccinate Tris Salt (CHS)** | Anatrace | CH210 | |
| **Benzonase® Nuclease HC, Purity > 90%** | Millipore Sigma | 71205 | 250 U/µl |
| **cOmplete™, Mini Protease Inhibitor Cocktail** | Roche (Millipore Sigma) | 11836153001 | |
| **Amicon Ultra Centrifugal Filter (MWCO, 100 kDa)** | Millipore Sigma | UFC910024 | |

#### Cell culture:

HEK293-GS22 cells were maintained in 75 $cm^2$, culture-treated, vented flasks (Corning) at in a humidified atmosphere at 37°C with 5% $CO_2$. Cell medium was 0.22 µm-filtered DMEM supplemented with 10% (v/v) FBS and penicillin/streptomycin. Cells were subcultured every 4-5 days at confluency.

HEK293FT cells were maintained under similar conditions except their medium was supplemented with 100-fold diluted 100X L-glutamine, 100X sodium pyruvate and, 100X MEM NEAA to working concentrations of 1X. Cultures were not tested for mycoplasma contamination.

#### Molecular biology:

Nanoluc (Nluc) was fused to the N-terminus of GLP-1R using Gibson assembly.[3] Briefly, plasmid DNA encoding Nanoluc (pNL1.1), provided by Promega, was reconstituted in 20 µL endotoxin free TE buffer (Qiagen). From the reconstituted plasmid template, the open reading frame of NanoLuc was linearized and amplified with the Phusion HF polymerase PCR kit (Thermo) according to the manufacturer's instructions. Human GLP-1R in a modified pSectag2 vector, kindly provided by Adam Schwaid (Pfizer), was linearized with appropriate overhangs to facilitate Gibson assembly with the Phusion HF polymerase PCR kit (Thermo) according to the manufacturer's instructions. The PCR products were treated with DpnI (0.8 µL, Aglient) for 1 h at 37 °C, and DNA was isolated using concentrator spin-columns (Zymo). 0.03 pmol and 0.06 pmol of linearized DNA encoding receptor + vector and Nanoluc, respectively, were assembled using 10 µL of 2X master mix at 50 °C for 1 h according to manufacturer's instructions (NEB). 5-alpha Competent *E. coli* (30 µL, NEB) were then transformed with 2 µL of assembly reaction mixture by heat shocking for 30 s at 42 °C and grown with SOC medium (0.5 mL) for 1 h at 37 °C with shaking. The cell suspension was spread onto LB agar plates with ampicillin (100 µg/mL), incubated overnight at 37 °C, and colonies were selected for plasmid Maxiprep according to the manufacturers protocol. Sanger sequencing performed at the UW-Madison Biotechnology Center DNA Sequencing Facility was used to confirm the desired sequence.

PCR primers (Overhangs shown in **bold**):

Nanoluc linearization forward primer:

5'-GGTATGGTCTTCACACTCGAAGATTTCGTTG-3'

Nanoluc linearization reverse primer:

5'- ACCACCCGCCAGAATGCGTTC-3'

Receptor + vector linearization forward primer

5'-**ATTCTGGCGGGTGGT**CCTCGCCCTCAGG-3'

Receptor + vector linearization reverse primer:

5'-**GTGTGAAGACCATACC**ATCCCCAGTGGAACCTGG-3'

Concatenated Sequencing reads:

```
ATGGAGACAGACACACTCCTGCTATGGGTACTGCTGCTCTGGGTTCCAGGTTCCACTGGGGATGGTATGG
TCTTCACACTCGAAGATTTCGTTGGGGACTGGCGACAGACAGCCGGCTACAACCTGGACCAAGTCCTTGA
ACAGGGAGGTGTGTCCAGTTTGTTTCAGAATCTCGGGGTGTCCGTAACTCCGATCCAAAGGATTGTCCTG
AGCGGTGAAAATGGGCTGAAGATCGACATCCATGTCATCATCCCGTATGAAGGTCTGAGCGGCGACCAAA
```

TGGGCCAGATCGAAAAAATTTTTAAGGTGGTGTACCCTGTGGATGATCATCACTTTAAGGTGATCCTGCA
CTATGGCACACTGGTAATCGACGGGGTTACGCCGAACATGATCGACTATTTCGGACGGCCGTATGAAGGC
ATCGCCGTGTTCGACGGCAAAAAGATCACTGTAACAGGGACCCTGTGGAACGGCAACAAAATTATCGACG
AGCGCCTGATCAACCCCGACGGCTCCCTGCTGTTCCGAGTAACCATCAACGGAGTGACCGGCTGGCGGCT
GTGCGAACGTATTCTGGCGGGTGGTCCTCGCCCTCAGGGCGCTACCGTGTCCCTCTGGGAAACCGTCCAG
AAGTGGAGGGAGTACCGTAGACAATGCCAGCGTAGCTTGACCGAGGACCCTCCTCCCGCGACAGACCTCT
TCTGTAACAGGACGTTCGATGAATACGCATGCTGGCCTGATGGAGAGCCGGGCTCATTCGTTAACGTCAG
CTGTCCGTGGTATCTGCCTTGGGCTTCCTCTGTTCCCCAGGGTCATGTCTACCGCTTCTGTACGGCAGAA
GGTCTGTGGCTCCAAAAAGACAACAGTTCGTTGCCTTGGCGCGACCTGTCCGAATGCGAGGAATCAAAAA
GAGGCGAACGCTCGTCTCCAGAGGAACAACTGCTGTTCCTTTACATTATCTACACGGTAGGATACGCCCT
TTCTTTCTCGGCGCTCGTAATCGCCTCGGCGATCCTGCTTGGCTTTCGCCACCTGCACTGCACTAGGAAC
TACATTCACCTGAATTTGTTCGCTTCATTCATCTTGCGTGCCCTCTCAGTTTTCATTAAGGACGCCGCTC
TTAAGTGGATGTACAGCACAGCTGCTCAACAGCATCAGTGGGACGGTCTGTTGTCCTACCAGGACAGTCT
GAGTTGTCGTCTGGTCTTCCTGCTGATGCAGTATTGTGTCGCAGCAAACTACTATTGGCTCTTGGTTGAG
GGTGTCTACCTGTACACTCTTTTGGCCTTCAGCGTGTTTTCTGAGCAGTGGATCTTCAGATTGTACGTTT
CCATCGGCTGGGGAGTGCCCCTTCTCTTTGTTGTGCCTTGGGGCATAGTTAAGTACCTTTACGAGGATGA
AGGTTGCTGGACAAGAAATTCTAATATGAACTACTGGCTGATTATCCGTCTGCCTATTCTCTTTGCCATT
GGTGTAAACTTTTTGATCTTCGTAAGGGTAATCTGCATAGTCGTATCTAAGTTGAAGGCTAACTTGATGT
GTAAAACCGATATCAAGTGTCGCCTGGCCAAGTCAACACTCACTTTGATCCCCCTTCTGGGTACGCACGA
GGTAATCTTTGCGTTTGTCATGGACGAACACGCCCGTGGTACTCTCCGCTTCATCAAGTTGTTTACCGAG
TTGTCCTTTACCAGTTTCCAAGGACTCATGGTGGCCATTCTGTACTGCTTCGTCAATAACGAGGTCCAGT
TGGAGTTCCGCAAGTCATGGGAACGCTGGCGCCTGGAACACCTGCATATCCAGCGCGACTCCTCTATGAA
ACCGCTGAAGTGTCCAACCAGTTCATTGTCATCCGGAGCTACTGCGGGATCTTCCATGTATACCGCCACT
TGTCAGGCTTCATGTTCTCTCGAGCATCATCACCACCACCACGACTATAAGGACGATGATGACAAGTAA

Translated Sequence (signal peptide = red; Nanoluc = blue; GLP-1R = green; linkers = black; affinity tags = purple):

METDTLLLWVLLLWVPGSTGDGMVFTLEDFVGDWRQTAGYNLDQVLEQGGVSSLFQNLGVSVTPIQRIVL
SGENGLKIDIHVIIPYEGLSGDQMGQIEKIFKVVYPVDDHHFKVILHYGTLVIDGVTPNMIDYFGRPYEG
IAVFDGKKITVTGTLWNGNKIIDERLINPDGSLLFRVTINGVTGWRLCERILAGGPRPQGATVSLWETVQ
KWREYRRQCQRSLTEDPPPATDLFCNRTFDEYACWPDGEPGSFVNVSCPWYLPWASSVPQGHVYRFCTAE
GLWLQKDNSSLPWRDLSECEESKRGERSSPEEQLLFLYIIYTVGYALSFSALVIASAILLGFRHLHCTRN
YIHLNLFASFILRALSVFIKDAALKWMYSTAAQQHQWDGLLSYQDSLSCRLVFLLMQYCVAANYYWLLVE
GVYLYTLLAFSVFSEQWIFRLYVSIGWGVPLLFVVPWGIVKYLYEDEGCWTRNSNMNYWLIIRLPILFAI
GVNFLIFVRVICIVVSKLKANLMCKTDIKCRLAKSTLTLIPLLGTHEVIFAFVMDEHARGTLRFIKLFTE
LSFTSFQGLMVAILYCFVNNEVQLEFRKSWERWRLEHLHIQRDSSMKPLKCPTSSLSSGATAGSSMYTAT
CQASCSLEHHHHHHDYKDDDDK*

#### Luminescence and BRET assays:

##### cAMP-Glosensor luminescence receptor activation assays:

The Glosensor protocol was adapted from Hager et al.[2] and Binkowski et al.[4] HEK293 cells stably expressing the Glosensor-22F (Promega) luminescent cAMP-sensing protein were grown to confluence, harvested, and then 1/3 of the collected cells were plated onto a 10 cm tissue-treated dish with 10 mL 10% FBS in DMEM without penicillin/streptomycin. The cells were then incubated at 37°C with 5% $CO_2$ overnight. After the overnight incubation, the medium was aspirated, 4.5 mL McCoy's 5A modified medium with 10% FBS was added, and the cells were incubated at 37°C with 5% $CO_2$. During this incubation, 5 µg of GLP-1R receptor plasmid (as a solution in endotoxin-free TE buffer (Qiagen); the construct contained a C-terminal FLAG tag and a $His_6$ tag immediately upstream of the FLAG tag) and 15 µL FuGENE HD transfection reagent was added to 1 mL of opti-MEM. After 20 min, 4.5 mL of DMEM with 10% FBS was added to the cells, and 1 mL of the transfection mixture was gently pipetted into the medium. The cells were then returned for incubation overnight. The next day, cells were washed with Dulbecco's phosphate buffered saline (DPBS), harvested with 0.05% Trypsin-EDTA, and resuspended into 4 mL 10% FBS in DMEM without penicillin/streptomycin. The cell suspension was diluted to approximately 500,000 cells/mL in medium, and 100 µL of cell suspension was pipetted into each well (providing ~50,000 cells/well) of a white-bottom, white-walled, 96-well plate. The cells in the 96-well plate were incubated at 37°C with 5% $CO_2$ overnight. After 24 h, the medium was removed by inverting and gently flicking the plate. DPBS with D-luciferin (500 µM) was quickly added to the plate (90 µL/well). The plate containing cells was allowed to sit for approximately 20 min at room temperature before addition of peptide (as 10 µL/well diluted in DPBS). Pipette tips were changed between serial dilutions. The plate was then transferred to a BioTek Synergy 2 plate reader with no optical filter ("hole"), 1 mm vertical probe offset, and read with a sensitivity value of 200. Curves were generated from luminescence values observed between 10 and 20 min.

Experiments were conducted with $n \geq 3$ biological replicates. Reported $EC_{50}$ and %Max values were a result of normalizing, averaging, and then fitting data to three-parameter sigmoidal curves in GraphPad Prism 6. The bottom of the curves was constrained to 0%. Normalization was performed with 100% representing the top of GLP-1's curve for individual experiments and 0% representing the luminescence value in absence of peptide.

#### GLP-1R cAMP inhibition assays:

HEK293 cells stably expressing the Glosensor-22F (Promega) luminescent cAMP-sensing protein were harvested, transfected, and transferred to 96 well plates identically as above (cAMP-activation assay) except using 10 µg of hGLP-1R plasmid (as a solution in endotoxin-free TE buffer (Qiagen), pCMV6-XL5 [hGLP1-R]) and 30 µL FuGENE HD reagent were used for the transfection step. Forty-eight hours after transfection, the medium was removed from the plates by inverting and gently flicking the plate. DPBS with D-luciferin (500 µM) was quickly added to the plate (90 µL/well). The plate containing cells was allowed to sit for approximately 5 min at room temperature before addition of peptides (as 10 µL/well diluted in DPBS). Pipette tips were changed between serial dilutions. The plate was then transferred to a BioTek Synergy 2 plate reader with no optical filter ("hole"), 1 mm vertical probe offset, and read with a sensitivity value of 200. After 15 min, GLP-1 (7-36)-$NH_2$ was added to each well to a final concentration of 250 pM and the plate was re-read for 30 min. Inhibition curves were generated from luminescence values observed between 10 and 20 min.

Experiments were repeated $n \geq 3$ biological replicates. Each experiment consisted of at least 7 different concentrations of peptide, with solutions prepared via serial dilution of a stock solution of each peptide prepared from a 1 mM stock in DMSO. For individual experiments, data from the selected timepoint were fit to 3-three-parameter sigmoidal curves with shared top values. Bottom values were constrained for GLP-1(*S,S*-ACPC) and Ex-4(*S,S*-ACPC) based on their maximal luminescence in the absence of GLP-1. Exendin 9-39 was constrained with a bottom value of 0%. Top and bottom values were applied to normalize the data to 100% and 0%, respectively. Then, the normalized data was averaged among the biological replicates and once again fit to a 3-three-parameter sigmoidal curve.

#### Bioluminescence resonance energy transfer (BRET) arrestin recruitment assay:

The protocol for arrestin recruitment assays was adapted from Hager et al.[2] and Jorgensen et al.[5] HEK293FT cells were grown confluence, harvested, and then 1/3 of the collected cells were plated onto a 10 cm tissue-treated dish with 10 mL 10% FBS in DMEM (with NEAA, L-glutamine, and sodium pyruvate, see above) without penicillin/streptomycin. The cells were then incubated at 37°C with 5% $CO_2$ overnight. After 24 h, a transfection mixture was made with 1:1 polyethylenimine (PEI, 1 mg/mL in water, pH 7.0):DNA in 1 mL opti-MEM. Either GFP$^2$-β-arrestin-1 (14 μg) or GFP$^2$-β-arrestin-2(R393E, R395E) (14 μg) along with GRK5 (250 ng) and GLP-1R-Rluc8 (250 ng for β-arrestin-1 experiments or 130 ng for β-arrestin2 experiments) was added to the PEI/Opti-MEM mixture, and the resulting transfection mixture was incubated for 20 min at room temperature. The cell medium was then aspirated, replaced with 4.5 mL of DMEM without FBS, and the transfection mixture was gently pipetted into the medium. Six hours after transfection (with incubation at 37°C with 5% $CO_2$), 4.5 mL of DMEM supplemented with 20% FBS was added to the dish. Twenty-four hours after transfection (with incubation at 37°C with 5% $CO_2$), cells were washed with DPBS, harvested with 0.05% Trypsin-EDTA, and resuspended into 4 mL 10% FBS in DMEM (with NEAA, L-glutamine, and sodium pyruvate, see above). The cell suspension was diluted to approximately 1,000,000 cells/mL in medium, and 100 µL of cell suspension was pipetted to each well (providing ~100,000 cells/well) of a white-bottom, white-walled, 96-well plate. The cells in the 96-well plate were incubated at 37°C with 5% $CO_2$ overnight.

Twenty-four hours after adding cells to the 96-well plate, the medium was removed by pipette, the cells were washed twice with DPBS (with glucose, 100 µL/well), and 100µL of DPBS (with glucose) was added to each well. The cells were incubated at 37°C with 5% $CO_2$ for 45 min to1 h before addition of peptide (as 10 µL dilutions in DPBS). After addition of peptides, the cells were allowed to sit at room temperature for 20 min before addition of the Rluc8 substrate, DeepBlueC (10 µL/well of 60 µM DeepBlueC in 2:1 DPBS:ethanol). The 96-well plate was transferred to a BioTek Synergy 2 plate reader with 400 nm (20 nm bandwidth) and 528 nm (30 nm bandwidth) optical filters, 1 mm vertical probe offset, 1-second integration time and read at maximum (200) sensitivity. Concentration-response curves were generated with $I_{528nm}/I_{400nm}$ values taken between 15 and 45min after initial read, as signal variability was found to be relatively high at earlier timepoints.

Experiments were repeated n ≥ 3 biological replicates. Each experiment consists of at least 7 different concentrations of peptide, with solutions prepared via serial dilution of a stock solution of each peptide prepared from 1 mM peptide stock in DMSO. Reported $EC_{50}$ and %Max values were a result of normalizing, averaging, and fitting data to three-parameter sigmoidal curves in GraphPad Prism 6. The bottom of the curves was constrained to 0%. Normalization was performed with 100% representing the top of GLP-1's curve for individual experiments and 0% representing the bottom value if the curves were fit with raw data and constrained to have a shared minimum.

#### Bioluminescence resonance energy transfer (BRET) G protein conformation assay:

Cell membrane were prepared as described previously.[6] Briefly, HEK293A cells stably express hGLP-1R were transiently transfected with Gαs-Nluc, Gβ1, and Gγ2-Venus at 1:1:1 ratio (20 µg total DNA/T175 flask). 24 hours post transfection, cells were harvested and homogenized with a polytron homogenizer at 4°C in membrane buffer (20mM BisTris pH7.4, 50mM NaCl, 1mM $MgCl_2$, 1×P8340 (protease inhibitor cocktail, Sigma). Cell homogenate was applied to a stepped sucrose gradient (60%, 40%, homogenate) and centrifuged at 22500 rpm for 2.5 hours at 4°C. The layer between 40% and homogenate were collected and diluted in membrane preparation buffer and centrifuged at 30,000 rpm for 20 min at 4°C. The final pellet was resuspend in 100uL membrane buffer, aliquoted and stored in -80°C. total protein concentration was determined using nanodrop.

For membrane based G protein conformation assays, 5 µg per well of cell membrane was incubated with furimazine (1:1,000 dilution from stock) in assay buffer (1× HBSS, 10 mM HEPES, 0.1% (w/v) BSA, 1× P8340 protease inhibitor cocktail, 1 mM DTT and 0.1 mM PMSF, pH 7.4). The GLP-1R-induced BRET signal between $G\alpha_s$-Nluc and Gγ-Venus was measured at 30 °C using a PHERAstar (BMG LabTech with Emission 1: 475-30nm, and Emission 2: 535-30nm). Baseline BRET measurements were taken for 2 min at 15 second interval before addition of vehicle or increasing concentration of the ligands, and the measurement was continued for a further 10 minutes before GTP was added (30uM) to induce G protein dissociation. Data were corrected for baseline and vehicle treated samples.

#### Bioluminescence resonance energy transfer (BRET) whole-cell competition ligand binding assay:

HEK293 cells stably expressing the Glosensor-22F (Promega) luminescent cAMP-sensing protein were harvested, transfected, and transferred to 96 well plates identically as above (cAMP-activation assay) except 1.2 µg of nluc-GLP-1R plasmid (as a solution in endotoxin-free TE buffer) and 5 µL FuGENE HD reagent were used for the transfection step. Forty-eight hours after transfection, the medium was removed by inverting and gently flicking the plate. A solution of DPBS, 0.1% BSA, and 0.02% $NaN_3$ was added to the wells (90 µL/well) and the plate was transferred to a cold room (4 °C) for 15 min. Dilutions of unlabeled, competitor peptides were made in DPBS in a polypropylene 96-well plate. The energy acceptor, GLP-1(7-35)-Lys((5)-6-tetramethylrhodamine)-$NH_2$, was added (10 µL, 2 µM in DPBS) to each solution of diluted competitor peptide and mixed thoroughly. The resulting mixtures of competitor peptides and energy acceptor peptide (final concentration of 20 nM) were pipetted to the wells containing cells. The 96-well plate was covered with aluminum foil and allowed to sit at 4°C. After 16-24 h, the plate was removed from the cold room and allowed to equilibrate at room temperature for 5 min. A solution of either H-Coelenterazine (2 mL DPBS + 100 µL of 1.4 mM H-Coelenterazine in Ethanol) was prepared and added to the 96-well plate (10 µL solution/well). The plate was then transferred to a BioTek Synergy 2 plate reader and read with a 1 s integration time over the course of 1 h with 460 nm (40 nm bandwidth) and 590 nm (35 nm bandwidth) optical filters with a sensitivity value of 170.

Concentration-response curves were generated with maximum $I_{590nm}/I_{460nm}$ values for each well over the experiment time course. Experiments were repeated with $n \geq 3$ biological replicates. Each experiment consisted of at least 7 different concentrations of peptide, with solutions prepared via serial dilution of a stock solution of each peptide prepared from a 1 mM peptide stock in DMSO. For individual experiments, the calculated BRET data (maximum $I_{590nm}/I_{460nm}$ values) were fit to 3-three-parameter sigmoidal curves with shared top and bottom values. Those top and bottom values were applied to normalize the data to 100% and 0%, respectively. Then, the normalized data was averaged among the biological replicates and once again fit to a 3-three-parameter sigmoidal curves with top and bottom constraints of 100% and 0%, respectively.

#### Nonlinear fitting:

Data were processed in Microsoft Excel and GraphPad Prism 6 software packages. Concentration-response data were normalized for each experiment in Prism (*vide supra* for normalization parameters). Averages and sample standard deviations were calculated in either Excel or prism. Processed data from Excel were returned to Prism for curve fitting. $EC_{50}$ or $IC_{50}$ and maximal-response values, as well as curves, shown in figures and tables are from 3-parameter sigmoidal fits.

We applied the Black-Leff operation model to quantify the transduction coefficients (log(τ (efficacy)/$K_A$ (functional affinity)) of each peptide in activating the GLP-1R. Each set of peptides was fit to the equation below (Eq 1.) in GraphPad Prism 6, and log(τ/$K_A$) values, were extracted (as mean and SEM). $E$ represents the system response given a set of parameters. $E_{max}$ is the maximal response, $[A]$ is the concentration of the agonist, $K_A$ is the association constant of the agonist for the receptor, and τ is the receptor density [$R_t$] divided by the intrinsic agonist efficacy $K_E$. Taking the difference of transduction coefficients between an agonist and the reference peptide (GLP-1(7-36)-$NH_2$) provides the normalized transduction coefficient, Δlog(τ/$K_A$). A bias-factor, or ΔΔlog(τ/$K_A$) value, is calculated by determining the difference between the normalized transduction coefficients for two signaling pathways.

(Eq. 1) $$E = \frac{E_{max}\tau[A]}{[A](1+\tau)+K_A}$$

Standard deviations for ΔΔlog(τ/$K_A$) values were estimated by propagating extracted standard-errors for log(τ/$K_A$) using equation 2. Where $\sigma_{prop}.$ represents propagated standard deviation, and $SE^2_{ref1}$ and $SE^2_{ref2}$ represent the standard errors of the reference ligand (GLP-1) in the first and second pathway being compared, respectively. $SE^2_{lig1}$ and $SE^2_{lig2}$ represent the standard errors of the test ligand in the first and second pathway being compared, respectively.

(Eq. 2) $$\sigma_{prop}. = \sqrt{SE^2_{ref1} + SE^2_{ref2} + SE^2_{lig1} + SE^2_{lig2}}$$

To determine whether a ΔΔlog(τ/$K_A$) value was statistically significant, ΔΔlog(τ/$K_A$) values were compared using one-way analysis of variance (ANOVA) with Bonferroni's post-test.

**Nanobody 35 Preparation:** Nanobody 35 was prepared as previously described.[7]

**Complex Purification:** GLP-1R complexes were purified in accordance with the protocol reported by Liang et al.[8] with some modifications. Briefly, insect cell-pellets overexpressing FLAG-GLP-1R-His, Gαs, G Gβ1 and Gγ2 from 1.25 L of culture (~30 g) were thawed and suspended in 80 mL of 30 mM HEPES, 50 mM NaCl, 2 mM $MgCl_2$ and 5 mM $CaCl_2$ (pH 7.4) supplemented with 2 µL benzonase and 2 x cOmplete Protease Inhibitor Cocktail tablets. GLP-1R ligand (peptide as 2.5 mM stocks in $H_2O$) was added to a final concentration of 10 µM. The mixture was stirred for 30 min. at room temperature. To the mixture was added apyrase (10 µL) and ~1 mg nanobody 35. The mixture was stirred for another 30 min. at room temperature. After stirring, 20 mL of detergent solution (5% LMNG and 0.3% CHS w/v in dd$H_2O$) and 1.4 mL 5 M NaCl was added to the cell suspension. The suspension was Dounce homogenized 5-times with a tight pestle. Then, 80 mL of 30 mM HEPES, 50 mM NaCl, 2 mM $MgCl_2$, 5 mM $CaCl_2$ (pH 7.4) and 0.8 mL of 5 M NaCl were added. The resulting mixture was stirred for 1 h at 4 °C. Insoluble debris were removed by centrifugation (30,000 g for 15 min.), and the supernatant was filtered with a glass fiber prefilter. Anti-FLAG affinity gel (~3 mL) was equilibrated in 30 mM HEPES, 50 mM NaCl, 2 mM $MgCl_2$ and 5 mM $CaCl_2$ (pH 7.4) and added to the filtered supernatant. The resulting mixture was placed on a rotator for 2 h at room temperature. The resin was then transferred to a glass column and washed with 100 mL wash buffer (20 mM HEPES, 100 mM NaCl, 2 mM $MgCl_2$, 5 mM $CaCl_2$, 2.5 µM GLP-1R ligand, and 0.01% LMNG + 0.0006% CHS). Crude complex was eluted with 25 mL elution buffer (20 mM HEPES, 100 mM NaCl, 2 mM $MgCl_2$, 0.2 mg/mL FLAG peptide, 2.5 µM GLP-1R ligand, 10 mM EGTA, and 0.01% LMNG + 0.0006% CHS, pH 7.4). TCEP (0.5 M) was added to the elution volume to a final concentration of 0.1 mM. The elution mixture was concentrated to a volume of ~0.5 mL with a 100 kDa MWCO centrifugal concentrator, and this solution was filtered with a 0.22 µm centrifugal filter before purification by size exclusion chromatography (SEC). The complex was resolved with on a Superdex 200 Increase 1G/300 GL column with 0.9 mL/min SEC running buffer (20 mM HEPES, 100 mM NaCl, 2 mM $MgCl_2$, 2.5 µM GLP-1R ligand, 0.1 mM TCEP and 0.01% LMNG + 0.0006% CHS, pH 7.4) SEC fractions containing complex were collected, pooled, concentrated to ~4 mg/mL, flash frozen in liquid nitrogen, and stored at -80 °C for further use. Small aliquots of SEC fractions were directly flash frozen for negative stain electron microscopy.

**SDS-PAGE and Western Blotting:** Samples were prepared for SDS-PAGE and western blotting with a 1:1:1 mixture of sample, 10% (w/v) sodium dodecyl sulfate (aq.), and Laemmli loading buffer containing 2-mercaptoethanol. Sample mixtures were not heated before loading onto gels. SDS-PAGE samples (10 µL) were loaded and run on Mini-PROTEAN TGX Precast 4-15% gels. For western blots, proteins were transferred (20 V, 12 h, 4 °C) onto PVDF membranes and blocked with 5% bovine serum albumin. Antibody solutions were applied for 1 h at room temperature.

**Negative-stain transmission electron microscopy:** Samples collected and flash frozen directly from SEC fractions were thawed and diluted to ~0.01-0.03 mg/mL with SEC buffer (without detergent). Immediately after dilution, 4 µL of sample was spotted on freshly glow-discharged (positive polarity, air chamber, 10 mA, 30 s) EM grids (carbon film on Cu, 300 mesh). After 60 s, excess sample was blotted away with Whatman filter paper. Uranyl formate (10 µL, 0.77% w/v aqueous solution) was applied to the grid and blotted three times. On the third application of uranyl formate, the solution was allowed to sit on the grid for 30 s before blotting. The grid was dried and loaded into a Talos L120C microscope. Micrographs were collected at 120 kV accelerating voltage, 73,000 magnification, and approximately -0.5 to -1.0 µm defocus values. Data were imported into and processed with RELION 3.1.

**Cryo-electron microscopy:** Samples (3 µL) were applied to acetone-prewashed, glow-discharged Ultrafoil R1.2/1.3 Au 300 mesh grids (Quantifoil GmbH, Großlöbichau, Germany) and flash frozen in liquid ethane using a Vitrobot Mark IV (Thermo Fisher Scientific, Waltham, MA, USA). Blot force was set to 19 and blot time set to 10 s. The Vitrobot sample chamber was set to 100% humidity and 4°C. Data were collected on a Titan Krios G3i microscope (Thermo Fisher Scientific, Waltham, MA, USA) operated at an accelerating voltage of 300 kV with a 100 µm objective aperture at an indicated magnification of 105 000× in nanoprobe EFTEM mode and a spot size of 5. A Gatan K3 direct electron detector positioned post a Gatan Quantum energy filter (Gatan, Pleasanton, CA, USA), operated in a CDS mode with a slit width of 25 eV was used to acquire dose-fractionated images. Movies were recorded as compressed TIFFs in normal-resolution mode yielding a physical pixel size of 0.83 Å/pixel with an exposure time of 5.011 s amounting to a total exposure of 49.8 $e^-/Å^2$ for at an exposure rate of 9.94 $e^-/ Å^2$/second that was fractionated into 71 subframes. The target defocus was set to −1.5 µm with 0.1 µm increments between holes. Beam-image shift was used to acquire data from 9 surrounding holes after which the stage was moved to the next collection area.

**Cryo-electron microscopy processing:** All processing was performed in RELION 3.1 beta[9] unless specified otherwise. 5805 movies were imported and motion-corrected with MotionCor2.[10] CTF estimation was performed with GCTF v1.06.[11] Particle picking was performed with crYOLO[12] which yielded 3,450,481 initial particle projections. Reference free 2D classification was performed and classes were manually selected providing 1,284,047 projections. Initial 3D classification with alignment was performed with a lowpass filtered (16 Å) reference map derived from a GLP-1R/G-protein structure. Two favorable classes were manually selected providing 671,599 particle projections. These particles were subjected to Bayesian polishing, CTF refinement, and 3D-auto refinement. This process provided a map with a global resolution of 2.32 Å (0.143 FSC, with detergent micelle and alpha-helical domain masked) corresponding to "conformer 1."

A mask excluding the density for the G protein, detergent micelle, and the distal region of the ECD was used to perform 3D classification without alignment (30 iterations, regularization parameter $\tau$ of 10) on the particles comprising conformer 1. One class was manually selected with 221,955 particles. These particles were subjected to auto-refinement in RELION 3.1 providing a map with a global resolution of 2.51 Å (0.143 FSC, with detergent micelle and alpha-helical domain masked) corresponding to "conformer 2."

**Atomic Modelling:** The structure of ExP5/GLP-1R/Gs (PDB: 6B3J)[8] was used as a template and rigidly fit into the cryo-EM density. COOT[13] v0.8.9.2 and ISOLDE[14] (implemented in UCSF ChimeraX v0.93[15]) were used to manually fit and refine the models. Automated real-space refinement and validation were performed with the PHENIX v1.18.2 software package.[16]

#### Peptide Synthesis and characterization:

##### A. Peptide Synthesis:

Polypeptides were generated by standard Fmoc-based solid-phase synthesis using a combination of automated and manual microwave-assisted methods.

NovaPEG Rink amide resin was swollen in DMF for 15 min and transferred to a Liberty blue automated peptide synthesis instrument. Couplings were performed with 2.5 mL of 0.2 M Fmoc-protected amino acid in DMF, 0.5 mL of 1 M oxyma, and 1 mL 1 M DIC with microwave irradiation at 75 °C for 3 min. Deprotections were performed with 20% piperidine in DMF with 0.1 M oxyma at 90 °C for 2 min. The instrument was programmed to synthesize the C-termini of the peptides on resin and halt before the 4$^{th}$ residue from the N-terminus.

The resin was then split and transferred to Torviq polypropylene syringes along with a Teflon coated magnetic stir-bar to complete the synthesis by manual, microwave-assisted methods. From this point onwards, diastereomers were synthesized and purified independently in a sequential fashion (rather than in parallel) to avoid misidentification. Fmoc-protected amino acids (100 µmol) were dissolved in 1 mL DMF (biotech. grade) with 37.5 mg/mL HATU. DIEA (35 µL) was added to this mixture, and the mixture was vortexed briefly and then incubated at room temperature for 2-3 min before addition to the resin/deprotected resin-bound polypeptide. Coupling was then performed with stirring and microwave irradiation to 70°C for either 4 min or 12 min. (We recommend that the residue, β-residues, proline residues, arginine residues, and β-branched residues are coupled for longer than 4 min under these conditions). Histidine residues were coupled for 12 min at 50 °C. Deprotections were performed by adding 4 mL 20% (v/v) piperidine in DMF with stirring and microwave irradiation to 80°C for 2 min. Resin was washed with at least 4 mL DMF (4x) after coupling and deprotection steps.

After the final deprotection step, the resin was washed with 4 mL DMF (4x), then 4 mL dichloromethane (4x), and then allowed to dry on an aspirator. Cleavage of polypeptides from resin was performed with 3 mL peptide of 92.5% trifluoracetic acid (TFA), 5% thioanisole, and 2.5% 1,2-ethanedithiol (v/v) at room temperature with either stirring or rocking for 4 h. TFA was blown off with a stream of $N_2$, and the crude peptide was precipitated with addition of 30 mL of cold diethyl ether. Crude peptide was then pelleted by centrifugation at 3.5k RPM for 5 min. The supernatant was poured off, and the crude peptide pellet was dried under a $N_2$ stream.

For HPLC purification, the crude peptide pellet was then dissolved in either dimethyl sulfoxide (DMSO, 1-2 mL) or 1:1 Acetonitrile:$H_2O$ (1-2 mL) for peptides containing methionine. 100-150 µL of peptide mixture was injected onto the HPLC column and eluted with 12 mL/min of a 10-60% acetonitrile (with 0.1% TFA v/v) gradient in filtered Nanopure water (with 0.1% TFA v/v) over 60 minutes. Fractions were collected with a UV 220 nm-triggered automated fraction collector.

Peptide fractions were pooled, frozen over dry ice, and lyophilized. Lyophilized peptides were then dissolved in 1-2 mL of water with minimal acetonitrile to promote solubility of particularly hydrophobic peptides. Purity of the peptide was determined by HPLC monitoring at 220 nm, and concentration was determined by UV-Vis absorption at 280 nm. Molar extinction coefficients were estimated by assigning 5500, 1490 and 125 $M^{-1}cm^{-1}$ units to each tryptophan, tyrosine, and cysteine residue, respectively, and summing those values. Peptides were then aliquoted into 1.5 mL polypropylene centrifuge tubes and lyophilized. Peptide aliquots were dissolved in DMSO to a concentration of 1 mM for later use and stored frozen with refrigeration.

#### B. Peptide Characterization:

Analytical HPLC: 10-90% MeCN over 5 minutes + 0.01% formic Acid. XBridge C18 2.1 x 50mm column 0.750 mL/min, 5 µL injections. The 220 nm absorption peak appearing with a retention time of ~5 min is an instrument artifact observed in all injections including a control injection of ultrapure water (see below).

GLP-1 :HAE**G**TFTSDVSSYLEGQAAKEFIAWLVKGR-$NH_2$

Peptide sequences: Peptides containing substitutions at their 4th residue, which is glycine in the native peptides (shown **bolded** in the sequences), are designated by identifying the unnatural residue in parentheses, where: (D-Ala) = D-Alanine, (RRX) = (*R*,*R*)-*trans*-2-aminocyclopentanecarboxylic acid, (L-Ala) = L-Alanine, (SSX) = (*S*,*S*)-*trans*-2-aminocyclopentanecarboxylic acid

GLP-1 (D-Ala)

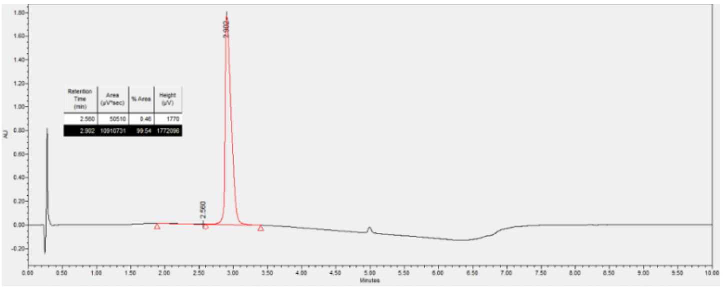

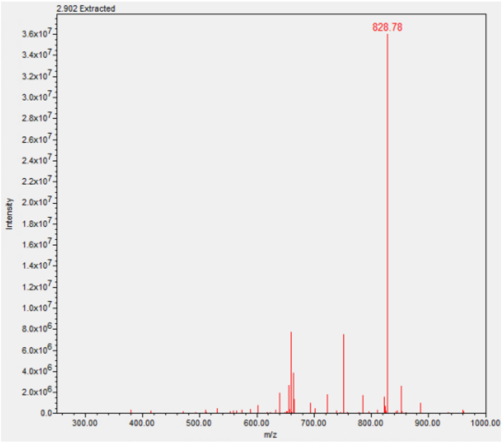

LCMS: >99%. For M = $C_{150}H_{228}N_{40}O_{45}$, calc. (most abundant *m/z*) $[M+4H]^{+4}$ = 828.6777, Found 828.78

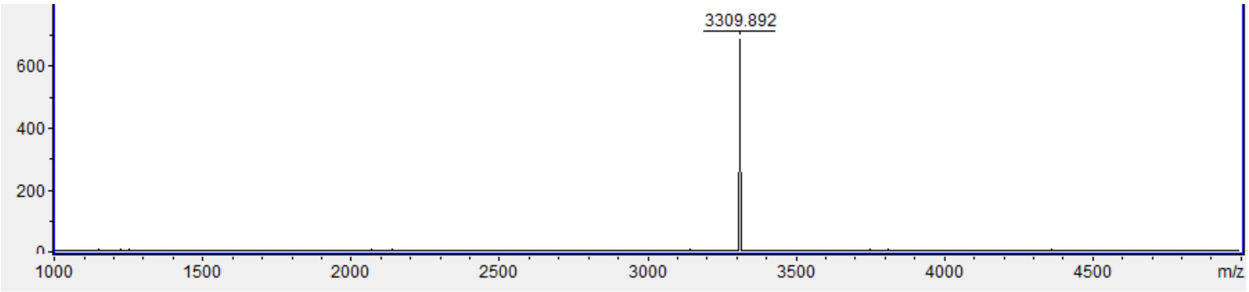

MALDI-TOF MS: For M = $C_{150}H_{228}N_{40}O_{45}$, calc. (monoisotopic *m/z*) $[M+H]^{+}$ = 3310.7, Found 3309.9.

GLP-1 (R,R-X)

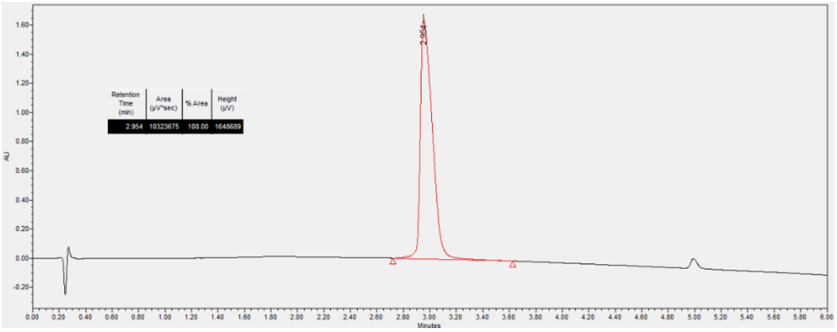

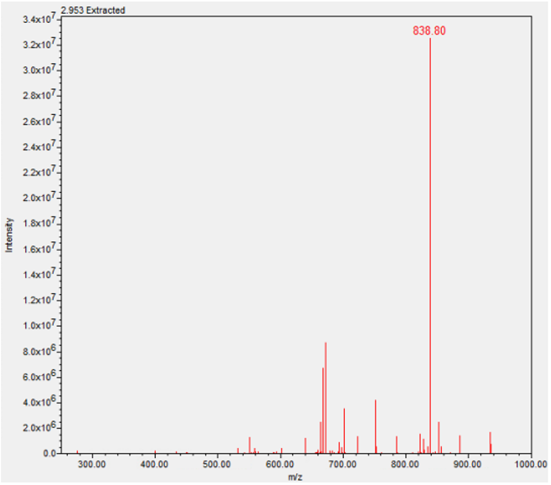

LCMS: >99%. For M = $C_{153}H_{232}N_{40}O_{45}$, calc. (most abundant *m/z*) $[M+4H]^{+4}$ 838.685, Found 838.80

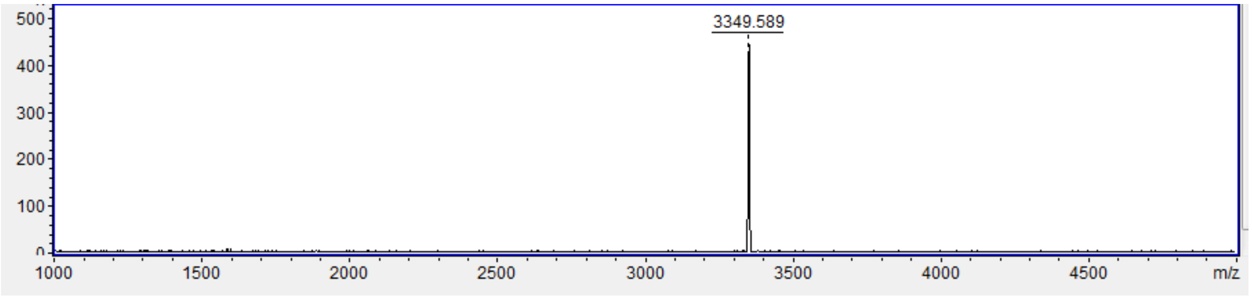

MALDI-TOF MS: For M = $C_{153}H_{232}N_{40}O_{45}$ calc. (monoisotopic *m/z*) $[M+H]^{+}$ = 3350.7, Found 3349.6.

GLP-1 (L-Ala)

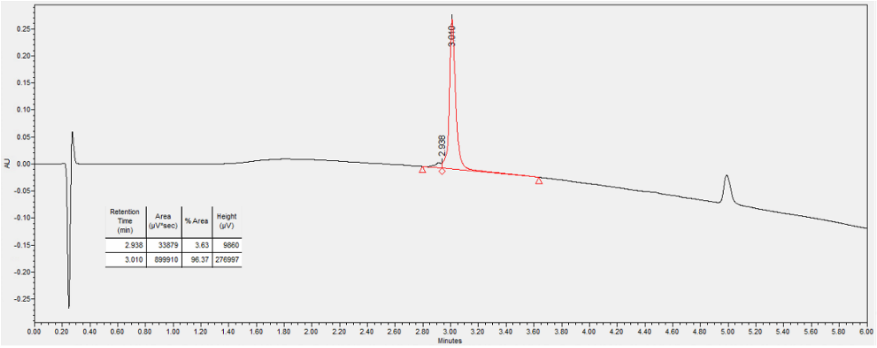

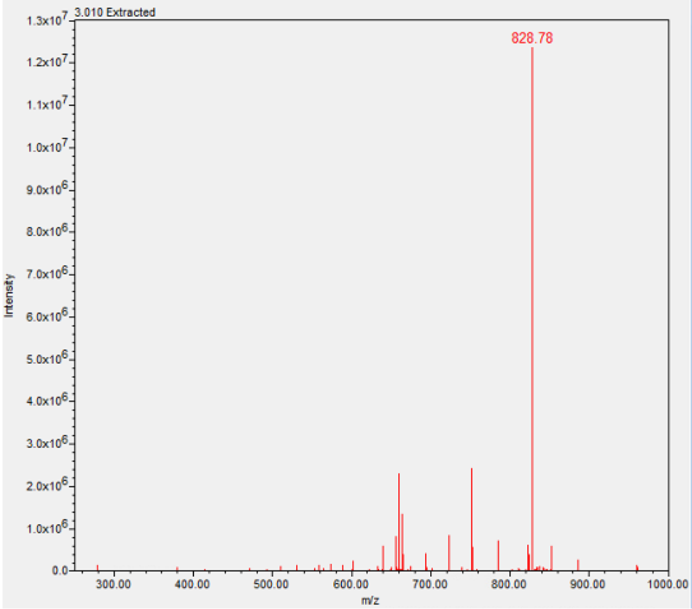

LCMS: 96%. For M = $C_{150}H_{228}N_{40}O_{45}$, calc. (most abundant *m/z*) $[M+4H]^{+4}$ = 828.677, Found 828.78

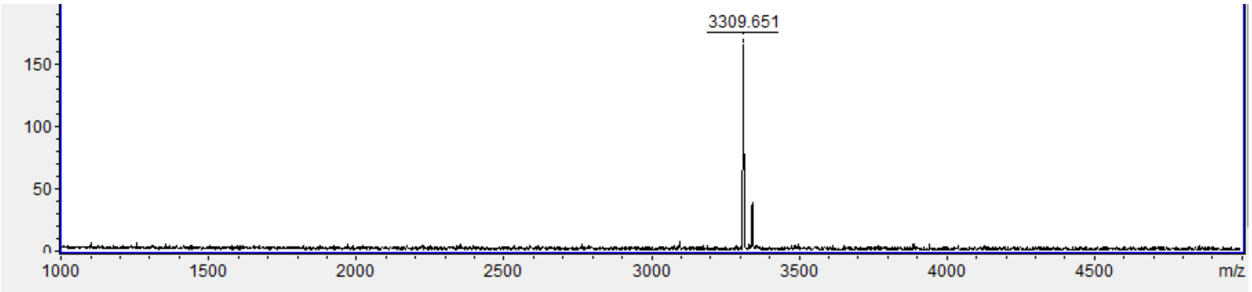

MALDI-TOF MS: For M = $C_{150}H_{228}N_{40}O_{45}$, calc. (monoisotopic *m/z*) $[M+H]^{+}$ = 3310.7, Found 3309.7.

GLP-1 (S,S-X)

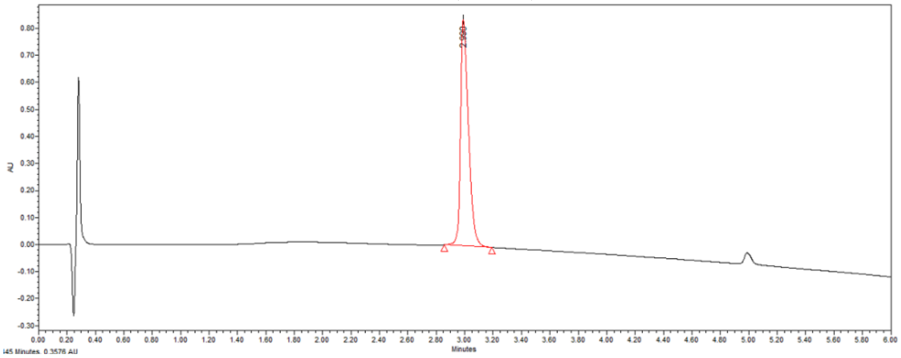

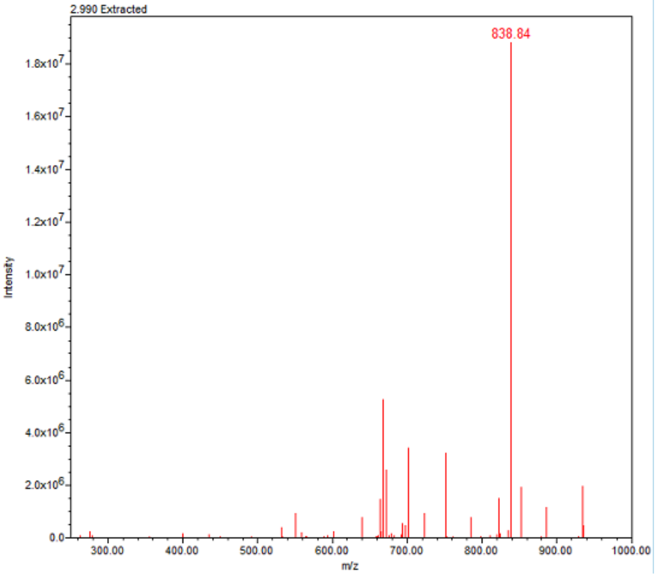

LCMS: >99%. For M = $C_{153}H_{232}N_{40}O_{45}$, calc. (most abundant *m/z*) $[M+4H]^{+4}$ = 838.685, Found 838.84

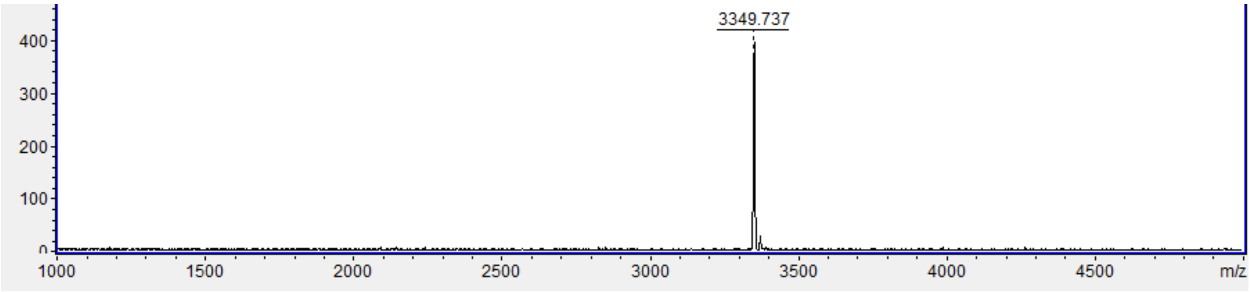

MALDI-TOF MS: For M = $C_{153}H_{232}N_{40}O_{45}$ Calc. (monoisotopic *m/z*) $[M+H]^{+}$ = 3350.7, Found 3349.7

Exendin-4 (D-Ala)

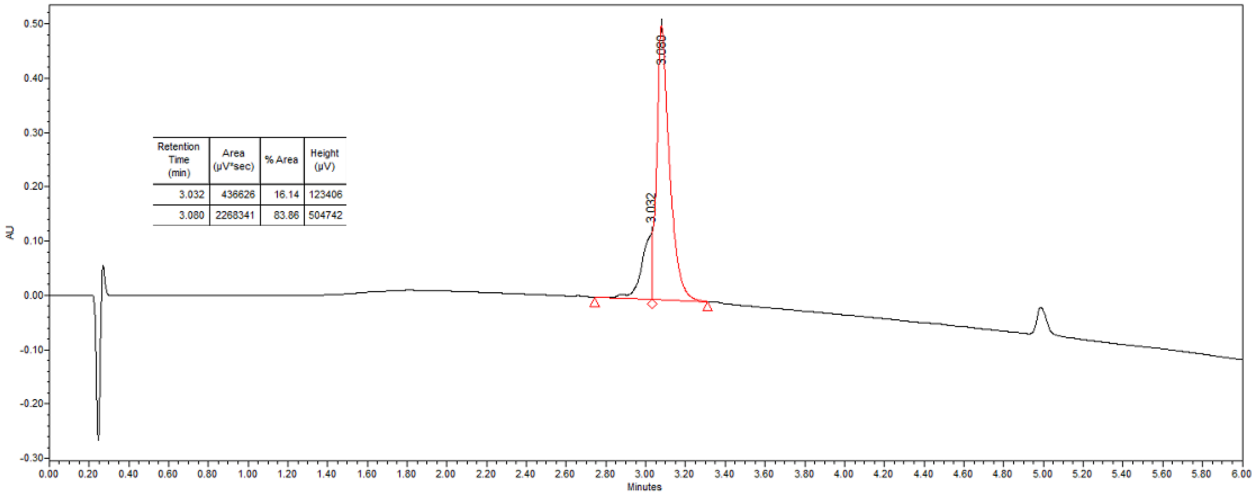

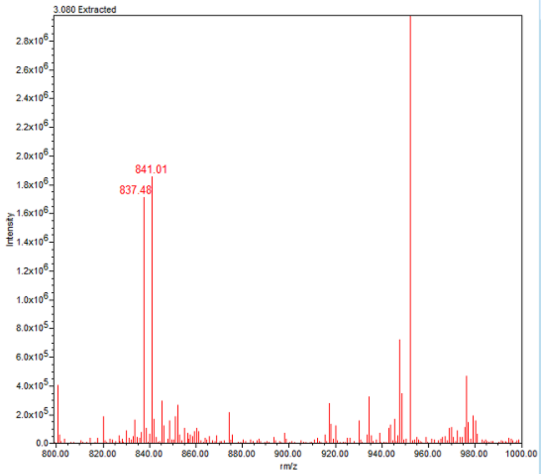

LCMS: 84%. For M = $C_{185}H_{284}N_{50}O_{60}S$, calc. (most abundant *m/z*) $[M+5H]^{+5}$=840.814. Found 841.01.

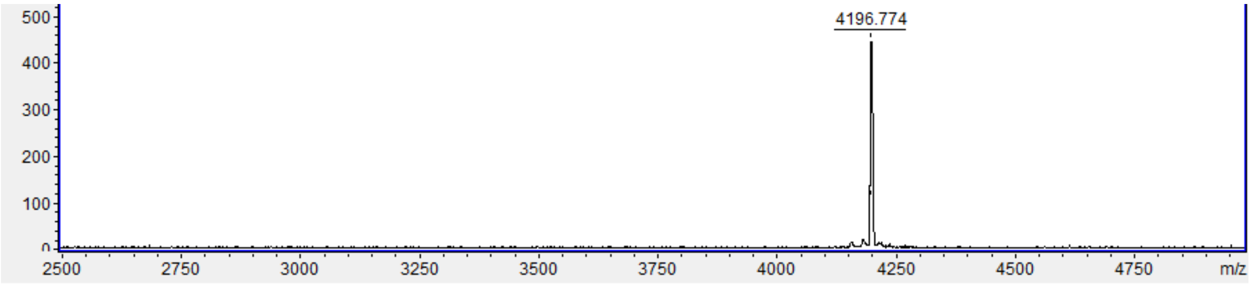

MALDI-TOF MS: For M = $C_{185}H_{284}N_{50}O_{60}S$, calc. (monoisotopic *m/z*) $[M+H]^{+}$ = 4198.0, Found 4196.7

Exendin-4 (R,R-X)

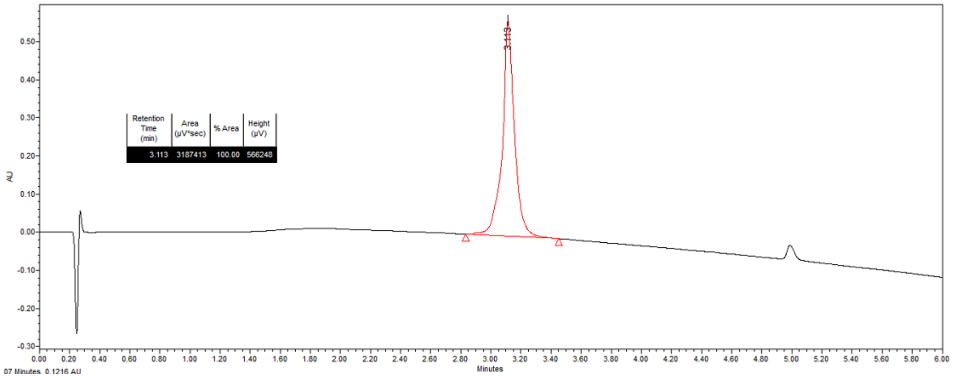

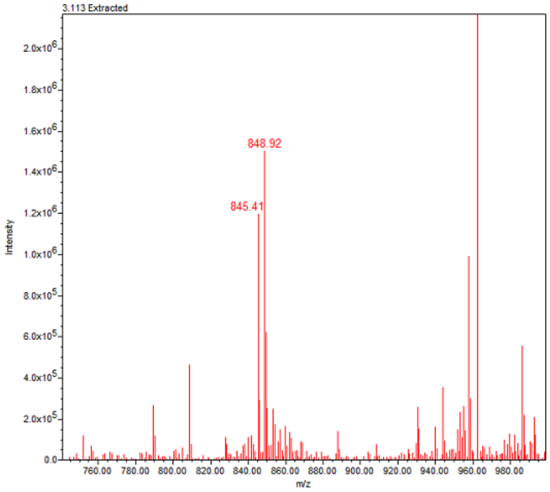

LCMS: >99%. For M= $C_{188}H_{288}N_{50}O_{60}S$, calc. (most abundant *m/z*) $[M+5H]^{+5}$ = 849.023. Found 848.92.

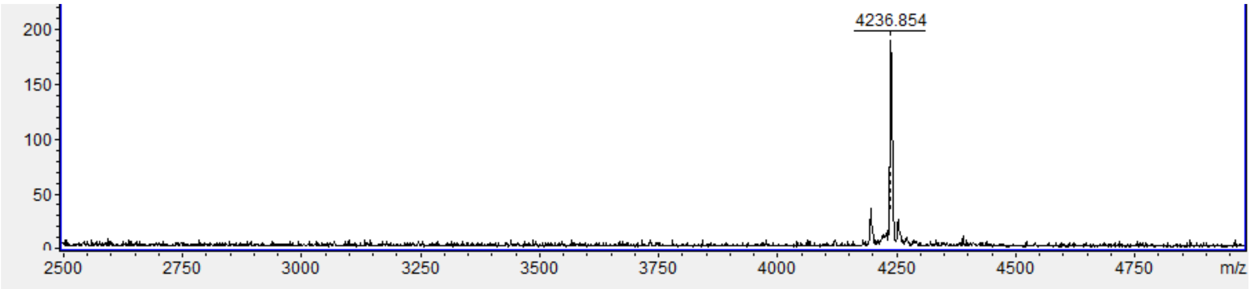

MALDI-TOF MS: For M= $C_{188}H_{288}N_{50}O_{60}S$, calc. (monoisotopic *m/z*) $[M+H]^{+}$ = 4239.1, Found 4236.8.

Exendin-4 (L-Ala)

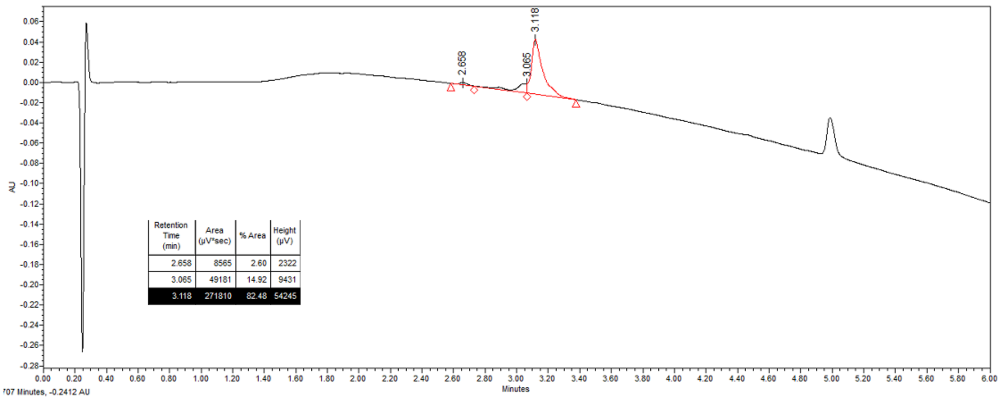

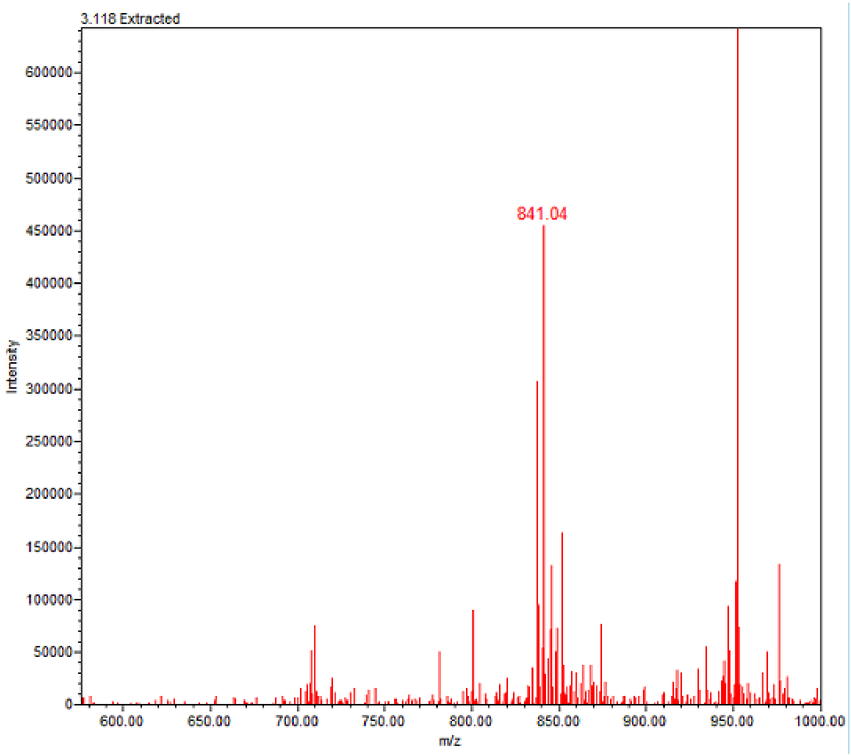

LCMS: 82%. For M = $C_{185}H_{284}N_{50}O_{60}S$, calc. (most abundant *m/z*) $[M+5H]^{+5}$=840.814. Found 841.04

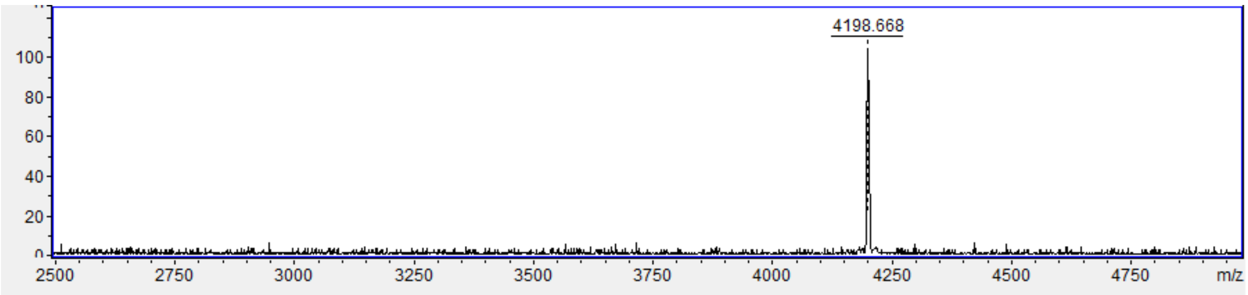

MALDI-TOF MS: For M = $C_{185}H_{284}N_{50}O_{60}S$, calc. (monoisotopic *m/z*) $[M+H]^{+}$ = 4198.0430, Found 4198.7.

Exendin-4 (S,S-X)

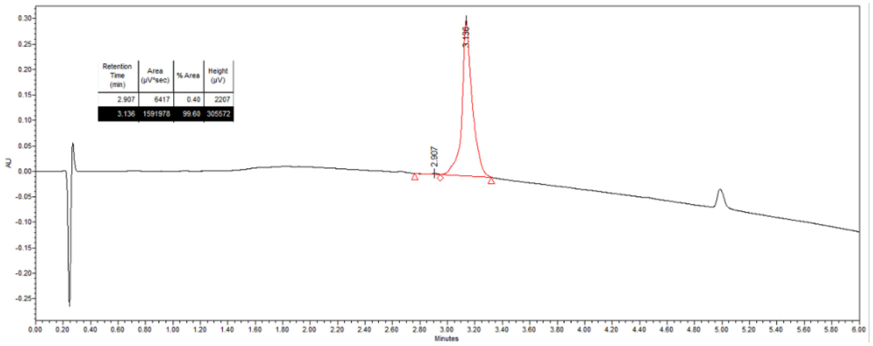

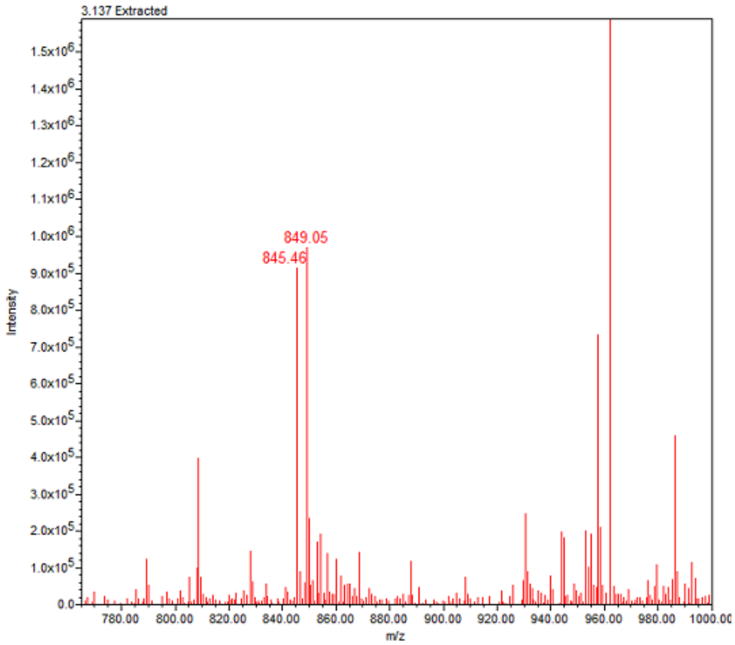

LCMS: >99%. For M= $C_{188}H_{288}N_{50}O_{60}S$, calc. (most abundant *m/z*) $[M+5H]^{+5}$=849.023. Found 849.05.

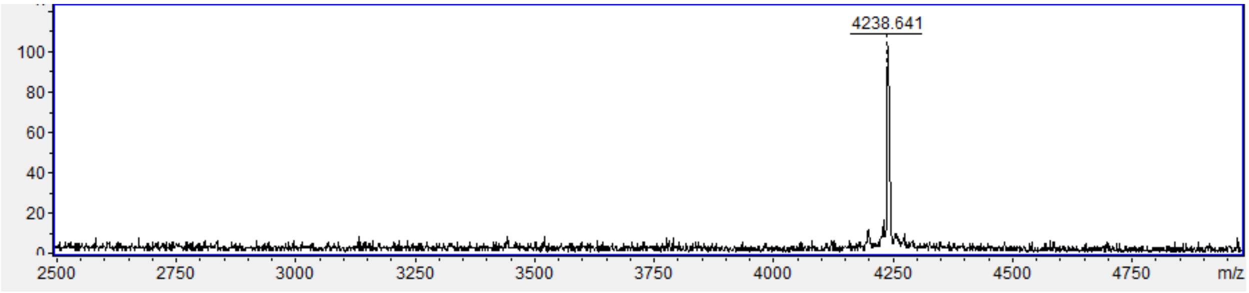

MALDI-TOF MS: For M= $C_{188}H_{288}N_{50}O_{60}S$, calc. (monoisotopic *m/z*) $[M+H]^{+}$ = 4239.0815, Found 4238.6.

5μL injection of Ultrapure $H_2O$

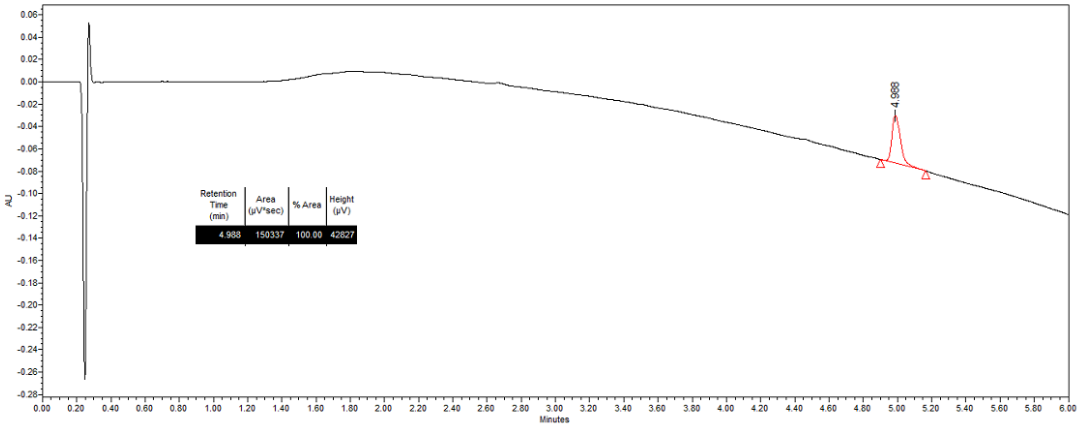
